# RNA-Encoded PGT121-LS Anti-HIV Antibody: Comprehensive Preclinical Characterization and Translational Pharmacokinetics

**DOI:** 10.64898/2026.06.24.734219

**Authors:** Felix Tolksdorf, Johannes Nelke, Robin Johannson, Janina Caesar, Anuhar Chaturvedi, Anja Kopp, Leyla Fischer, Alexandra Malz, Sven Kratochvil, Imke Gerhard, Jan P. Bogen, Candice Morin, Maximilian Kullmann, Michael S. Seaman, Georgia D. Tomaras, Nicole L. Yates, Margaret E. Ackerman, Joshua A. Weiner, Ursula Ellinghaus, Christiane R. Stadler, Ugur Sahin, Valentin Le Douce

## Abstract

Human Immunodeficiency Virus (HIV)-1 broadly neutralizing antibodies (bNAbs) have demonstrated clinical efficacy, but face manufacturing challenges associated with recombinant protein production and purification. Here, we present a ribonucleic acid (RNA)-encoded bNAb (RibobNAb) platform that enables *in vivo* antibody production of the clinically validated bNAb PGT121 via lipid nanoparticle (LNP) delivery, supporting rapid evaluation of Fc variants (LS, del294, LS-del294) *in vitro* and *in vivo*.

We confirmed expression, sub-nanomolar HIV-1 Env binding, and potent neutralization across all RibobNAb variants *in vitro*. In mice, single RNA-LNP administrations yielded *in vivo* expression of all RibobNAb variants, with PGT121-LS exhibiting a prolonged half-life compared with PGT121. In non-human primates (NHPs), a single intravenous administration of PGT121-LS RNA-LNP was well tolerated without anti-drug antibody (ADA) formation over 180 days and resulted in PGT121-LS half-lives comparable to the reference protein. Single intramuscular administration showed RibobNAb expression but resulted in ADA development from Day 14 onwards and lower bioavailability. *In vivo*-expressed PGT121-LS RibobNAb retained identical antiviral functionality to PGT121-LS reference protein. An NHP pharmacokinetics model integrating RNA transfection and translation dynamics enabled allometric scaling and first-in-human dose prediction. We highlight RibobNAbs as an alternative to conventional purified protein antibodies for rapid development of bNAb-based therapeutic strategies.

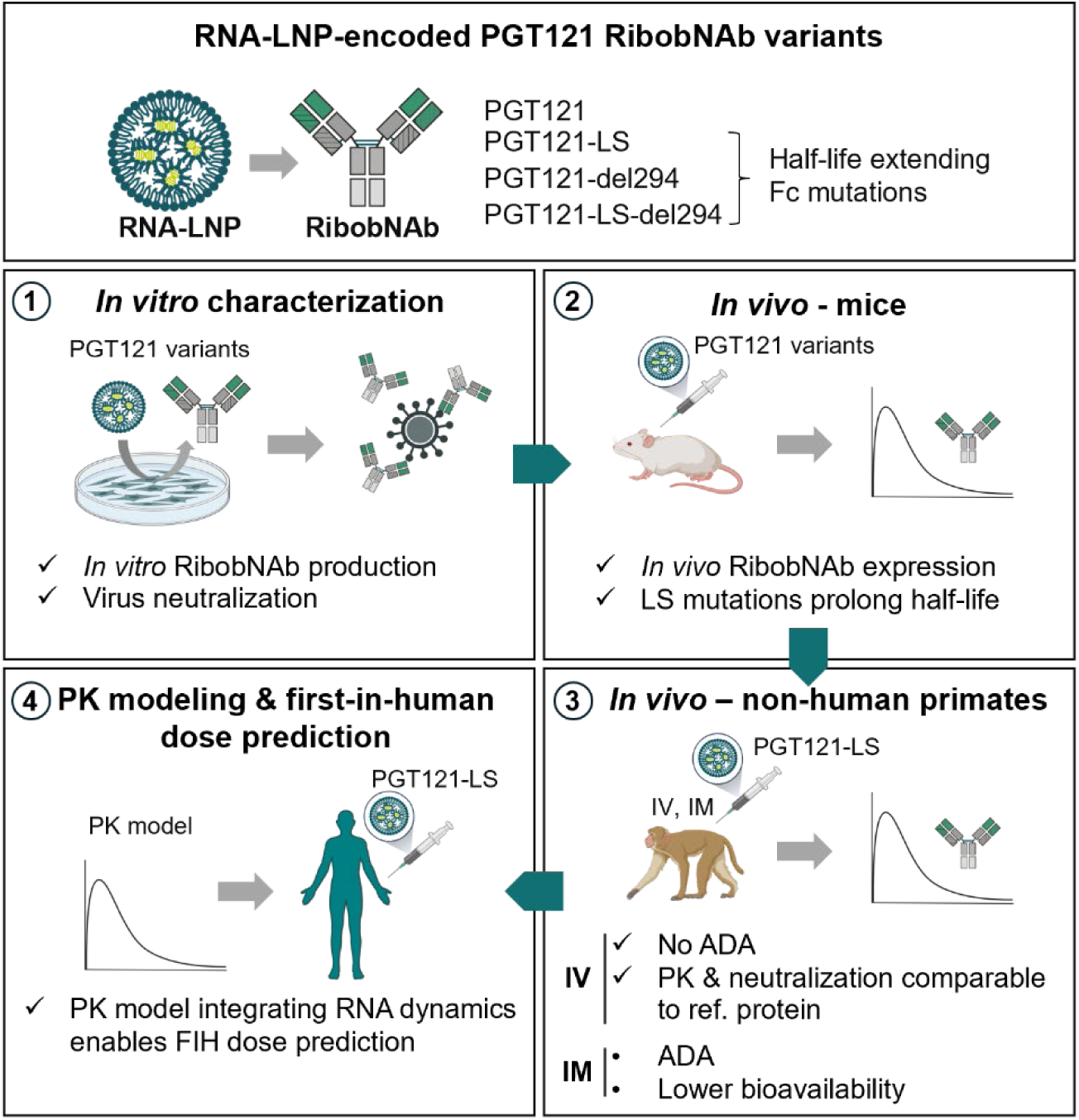
Graphical abstract.

## Introduction

Human Immunodeficiency Virus (HIV) remains a major global public health challenge, with approximately 40.8 million people infected worldwide.^1^ Although combined antiretroviral therapy (cART) has greatly improved HIV management by suppressing viral replication, challenges such as drug resistance, side effects, and adherence to strict daily regimens underscore the need for alternative therapeutic strategies enabling durable viral suppression in the absence of cART.^2^

Broadly neutralizing antibodies (bNAbs) have emerged as an alternative therapeutic approach. By targeting conserved regions of the HIV-1 envelope glycoprotein (Env), bNAbs confer cross-clade neutralization and offer both therapeutic and prophylactic potential.^3^ However, administration of bNAbs as purified proteins is often limited by production costs and scalability issues, particularly when large therapeutic doses are needed.^4,5^

Ribonucleic acid (RNA)-based therapeutics for *in vivo* protein production offer an alternative to purified protein delivery. Here, we have extended our previously reported RNA-encoded antibody platform developed for oncology^6–8^ to express RNA-encoded bNAbs (RibobNAbs). We anticipate that such RNA-encoded therapeutics will reduce manufacturing complexity and costs by eliminating large-scale cell culture, purification, and protein formulation processes,^9^ while also enabling rapid testing of diverse antibody formats. These include bispecific antibodies, antibody-like structures, fragment crystallizable (Fc) variants with modified effector functions or half-life, and alternative isotypes.^6,7,10–12^ *In vivo* antibody production can also provide native post-translational modifications and may reduce anti-drug antibody (ADA) formation, potentially improving immunogenicity profiles.^13^ In addition, the rapid clearance observed with some antibodies delivered as proteins is replaced by transient local production with sustained antibody levels, which may support more flexible administration schedules (e.g., fewer doses, intramuscular [IM] or subcutaneous administration [SC]).^9,14^

For *in vivo* delivery, RibobNAb-encoding RNA sequences are packaged into lipid nanoparticles (LNP) which, upon intravenous (IV) administration, are taken up by the liver, where the RNA is translated into polypeptide chains that self-assemble into functional bNAbs for secretion into the bloodstream.^6–8^

In this study, we generated RibobNAb variants of PGT121, a highly potent bNAb targeting an epitope on the V3 loop of glycoprotein 120 (gp120), including the conserved N332 glycan and adjacent glycans.^15,16^ In a phase 1 clinical trial (NCT02960581), a single IV infusion of PGT121 at 30 mg/kg was well tolerated and suppressed HIV RNA replication in viremic individuals, making it a potential component of for bNAb-based combination strategies.^17–20^

We performed preclinical studies to assess expression, binding, and neutralization potency of several PGT121 RibobNAb variants and selected PGT121-LS RibobNAb as our lead candidate. Non-human primate (NHP) pharmacokinetic (PK) studies enabled development of an RNA-encoded antibody PK model and first-in-human (FIH) dose predictions. This study describes the preclinical pharmacology of PGT121-LS RibobNAb, demonstrating the platform’s adaptability and potential to address challenges associated with recombinant bNAb therapeutics.

## Results

### PGT121 RibobNAb IgG1 variants are expressed and assembled following RNA or RNA-LNP transfection *in vitro*

The PGT121 RibobNAb variants evaluated in this study are encoded by two distinct nucleoside-modified RNA molecules encapsulated in LNPs (RNA-LNPs) previously designed and optimized for IV administration and subsequent translation in the liver (Figure 1A).^7,21,22^ The RNA molecules encode the light and heavy polypeptide chains of the PGT121 RibobNAbs, which self-assemble into functional antibodies consisting of two fragment antigen-binding (Fab) regions that target the V3 glycan epitope on HIV-1 Env and an IgG1 Fc region.^15,16^

**Figure 1:**
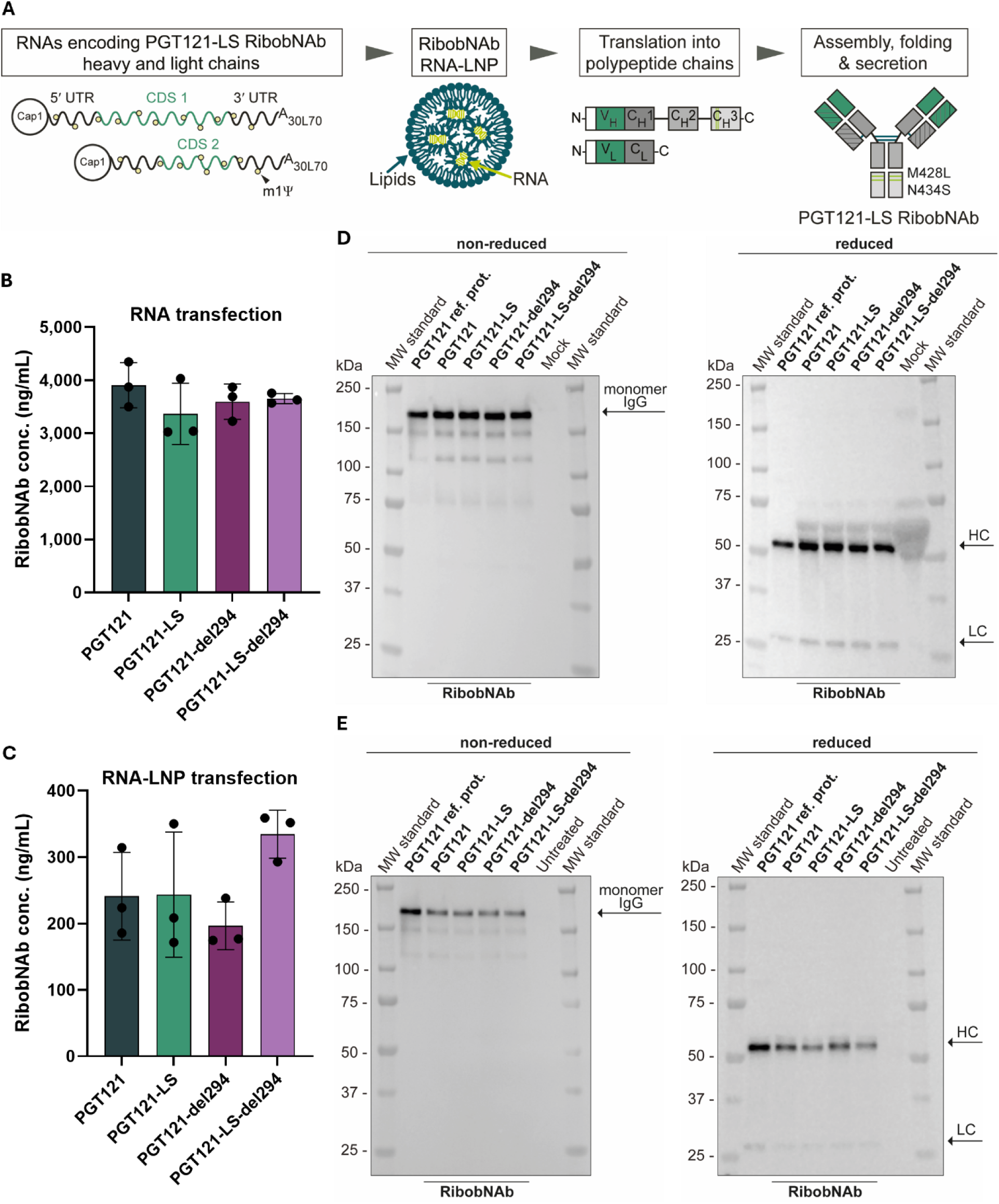
The RibobNAb platform enables efficient expression and correct assembly of PGT121 RibobNAb monomers. A) The translation of PGT121-encoding nucleoside-modified RNA into functional monomeric IgG antibodies is schematically shown for the PGT121-LS RibobNAb variant. Two RNA chains encoding the light and heavy chain of the antibody (first image) are encapsulated by an LNP formulation into RNA-LNPs (second image). Following *in vitro* transfection or *in vivo* administration, target cells translate these RNAs into the two single polypeptide chains of the antibody (third image) which are subsequently assembled and folded into the active monomeric IgG antibody (fourth image). The assembled RibobNAb is then secreted into the supernatant or circulation. The LS mutations (M428L/N434S) incorporated into the Fc region aim to enhance the pharmacokinetic properties of the antibody by improving binding to hFcRn at pH6. B, C) RibobNAb concentrations in HEK-293T-17 supernatant 48 h post RNA transfection (B) or RNA-LNP transfection (C) measured by ELISA. Error bars indicate the standard deviation (SD) of the mean of triplicates. D) Western blots of RibobNAb-containing HEK-293T-17 supernatants 48 h post RNA transfection. Each lane was loaded with samples containing 7 ng of protein. PGT121 reference protein served as positive control, while RiboJuice™-treated samples served as mock control. Samples were analyzed under non-reducing (left) and reducing conditions (right). E) Western blots of RibobNAb-containing HEK-293T-17 supernatants 48 h post RNA-LNP transfection. Each lane was loaded with samples containing 2.23 ng of protein. PGT121 reference protein and untreated samples served as controls. Samples were analyzed under non-reducing (left) and reducing conditions (right). Conc. = concentration; HC = heavy chain; IgG = immunoglobulin G; LC = light chain; LNP = lipid nanoparticle; ref. prot. = reference protein; RibobNAb = RNA-encoded, LNP-formulated anti-HIV-1 broadly neutralizing antibody; RNA = ribonucleic acid.

Leveraging the adaptability of the RibobNAb platform, we generated several PGT121 variants incorporating Fc modifications previously described to improve PK, while retaining the original Fab region.^23,24^ With the exception of the Fc mutations described below, the PGT121 RibobNAb variants are identical to the PGT121 reference protein in amino acid sequence.^15^

The PGT121 RibobNAb variants included a candidate with an unmodified Fc region (PGT121 RibobNAb), and candidates engineered with LS (M428L/N434S) mutations (PGT121-LS RibobNAb), an E294 deletion (PGT121-del294 RibobNAb), or a combination of both (PGT121-LS-del294 RibobNAb). The LS mutations improve antibody binding to the human neonatal Fc receptor (hFcRn), a receptor critical for antibody recycling, whereas the E294 deletion promotes hypersialylation of N297 in the Fc region, each prolonging antibody half-life through distinct mechanisms..^23–26^

We first confirmed RNA-mediated RibobNAb expression *in vitro*. HEK-293T-17 cells were transfected with 1 µg of RNAs encoding the heavy and the light chains, and RibobNAb levels in the cell culture supernatant were quantified using an enzyme-linked immunosorbent assay (ELISA). All RibobNAb variants were expressed at comparable levels, with concentrations ranging from 3,000 to 4,300 ng/mL (Figure 1B). To confirm that the detected RibobNAbs were correctly assembled into a native monomeric IgG quaternary structure, we assessed all variants by Western blot analysis. Under non-reducing conditions, a monomeric band at ∼150 kDa was observed for all variants, indicating correct assembly. Additionally, no visible aggregation or high molecular weight species were detected, and, under reducing conditions, distinct bands corresponding to the antibody heavy and light chains were observed (Figure 1D).

We then progressed to RNA-LNP transfection. For this method, HEK-293T-17 cells were transfected with 1 µg/mL of RNA-LNPs, yielding RibobNAb concentrations between 170 and 360 ng/mL (Figure 1C). Importantly, these values matched expression levels typically observed for RNA-encoded antibodies produced with the RiboMab platform.^7^ Western blot analysis confirmed that RibobNAbs generated via RNA-LNP transfection exhibited the same correct monomeric IgG assembly and chain composition as observed following RNA transfection (Figure 1E).

Together, these results confirm that our RNA-encoded antibody platform allows for efficient production of fully assembled antibodies and enables rapid screening of a panel of modified candidates. Importantly, modifications to the Fc region, including the LS mutations and del294 deletion, did not affect the expression or assembly of the RibobNAbs.

### PGT121 RibobNAb variants bind to native HIV-1 Env trimer with high affinity and retain neutralization potency *in vitro*

To confirm the functionality of *in vitro*-expressed RibobNAb variants, we assessed target binding and neutralization activity. Binding affinities of the RibobNAbs to the BG505 DS-SOSIP.664 Env Trimer (SOSIP),^27,28^ a stabilized mimetic of the HIV-1 Env trimer, were analyzed using bio-layer interferometry (BLI). Multi-cycle kinetic analyses were used to determine the association (k_on_) and dissociation (k_off_) rate constants, as well as the equilibrium dissociation constants (*K*_D_) (Figure 2A). All RibobNAb variants demonstrated high binding affinities to SOSIP, comparable to that of the PGT121 reference protein, with K_D_ values in the sub-nanomolar range (K_D_ RibobNAbs: 0.10–0.32 nM, K_D_ PGT121 reference protein: 0.81 nM) (Figure 2B). These values are consistent with the previously determined *K*_D_ value of 0.76 nM determined for the PGT121 reference protein, as measured by SPR.^29^

**Figure 2:**
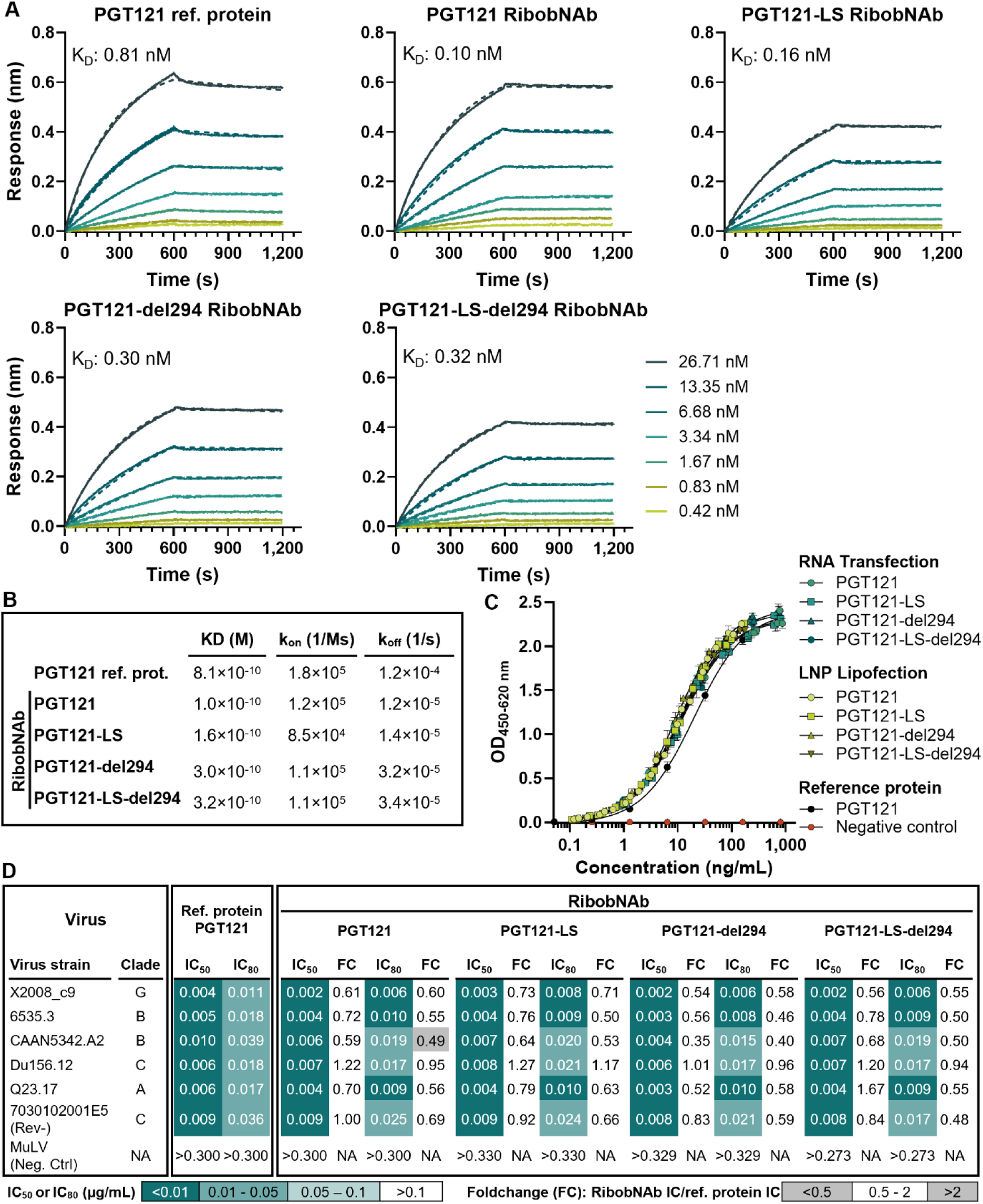
*In vitro*-expressed PGT121 RibobNAb variants bind to SOSIP with high affinities and neutralize a broad panel of HIV-1 strains. A) BLI sensorgrams showing binding kinetics of PGT121 reference protein and PGT121 RibobNAbs to SOSIP. The *in vitro*-expressed RibobNAbs (produced using RNA transfection) were immobilized on anti-hIgG Fc (AHC) biosensors. Binding affinities were determined using multi-cycle kinetics and decreasing SOSIP concentrations (26.71 – 0.42 nM, 2-fold dilutions). The association step was carried out for 600 s, followed by a 600 s dissociation step. B) Binding parameters determined by BLI. C) SOSIP-ELISA comparing the binding of *in vitro*-expressed PGT121 RibobNAb variants produced by either RNA transfection or RNA-LNP transfection to SOSIP. PGT121 reference protein served as positive control and an irrelevant antibody as negative control. Error bars indicate the SD of the mean of triplicates. D) TZM-bl pseudovirus neutralization assay against a diverse panel of viral strains using a dilution series of PGT121 recombinant protein or *in vitro* produced PGT121 RibobNAb variants. Foldchange (FC) values compare RibobNAb potency relative to recombinant protein. Values >0.5 and <2-fold indicate comparable activity. FC = foldchange; IC_50_ = half maximal inhibitory concentration; IC_80_ = 80% inhibitory concentration; LNP = lipid nanoparticle; ref. prot. = reference protein; RibobNAb = RNA-encoded, LNP-formulated anti-HIV-1 broadly neutralizing antibody; RNA = ribonucleic acid.

ELISA analysis confirmed that all PGT121 RibobNAb variants exhibited dose-dependent binding to SOSIP directly comparable to that of the PGT121 protein reference (Figure 2C).

To confirm the functionality of the RibobNAb variants, we evaluated their neutralization activity in a TZM-bl cell pseudovirus assay (Figure 2D) against a 6-Env panel of HIV-1 strains with known PGT121-sensitivity and representing different clades (A, B, C and G). None of the tested antibodies neutralized the negative control strain MuLV, confirming assay specificity. All RibobNAb variants effectively neutralized the PGT121-sensitive HIV-1 strains, with inhibitory concentrations (IC_50_ and IC_80_ values) within a two-fold range of the PGT121 reference protein for most HIV-1 strains, indicating equivalent neutralization efficacy. Neutralization potency was consistent across clades.

These results confirm that all RibobNAbs exhibit high binding affinities and neutralization activity comparable to the PGT121 reference protein.

### PGT121 RibobNAb variants are expressed *in vivo*, with LS mutations demonstrating enhanced PK in NSG Tg32 mice

After confirming expression *in vitro*, we proceeded to evaluate the *in vivo* expression and PK characteristics of the RibobNAbs in two murine models.

*In vivo* expression was first assessed in NSG mice. Mice received a single IV dose of 30 µg RNA-LNPs encoding either PGT121, PGT121-LS, PGT121-del294 or PGT121-LS-del294 RibobNAb (Figure 3A). Blood samples were collected over a 24-day period from subgroups at alternating time points (Figure 3A). All RibobNAbs were successfully expressed, achieving peak serum concentrations (C_max_) ranging from 400 to 600 µg/mL within 24–48 hours (h) post-administration and remained detectable throughout the study period.

**Figure 3:**
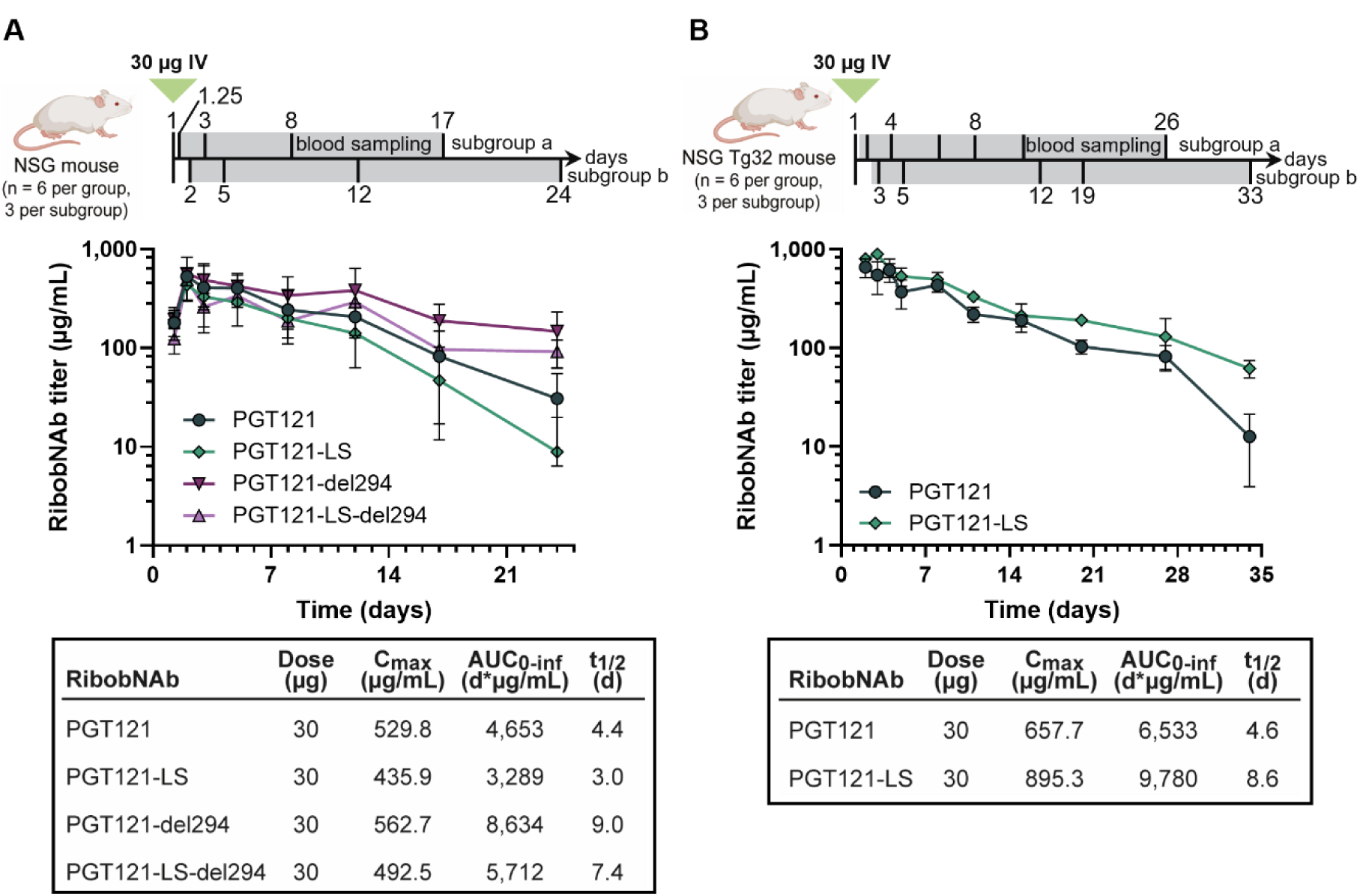
PGT121 RibobNAb variants are expressed in NSG mice and PGT121-LS shows improved pharmacokinetics in NSG Tg32 mice. A) PK analysis of PGT121 RibobNAb variants in NSG mice. Male and female NSG mice (n = 6 per group, 3 per subgroup) received a single IV bolus of 30 μg RNA-LNPs encoding either PGT121, PGT121-LS, PGT121-del294 or PGT121-LS-del294 RibobNAb. Blood samples were collected at the indicated time points using an alternating subgroup sampling strategy. B) PK analysis in NSG Tg32 mice. Male and female NSG Tg32 mice (n = 6 per group, 3 per subgroup) expressing hFcRn received a single IV bolus of 30 µg RNA-LNPs encoding either PGT121 or PGT121-LS RibobNAb. Blood samples were collected at the indicated time points using an alternating subgroup sampling strategy. A and B) RibobNAb serum concentrations were determined using ELISA. PK parameters were determined using NCA. Error bars indicate the SD of the mean of three animals. AUC_0-inf_ = Area under the serum concentration-time curve from time zero to infinity; C_max_ = maximum serum concentration; hFcRn = human neonatal Fc receptor; IV = intravenous; NSG = NOD scid gamma; NSG Tg32 = NOD scid gamma mice expressing hFcRn; RibobNAb = RNA-encoded, LNP-formulated anti-HIV-1 broadly neutralizing antibody; t_1/2_ = elimination half-life.

Building on these results, we next evaluated the impact of half-life extending mutations. The del294 deletion has been shown to increase antibody half-life in mice, but it abolishes Fc effector functions by disrupting Fc-gamma receptor (FcγR) binding and has not yet been tested in clinical settings.^23^ In contrast, the LS mutations retain Fc effector functions, which contribute substantially to *in vivo* antiviral activity, and have already been clinically validated.^30,31^ We therefore focused our subsequent studies on PGT121-LS RibobNAb. LS mutations enhance binding to hFcRn, thereby increasing antibody recycling.^30,31^ As the LS mutations do not impact binding to mFcRn, we used the transgenic NSG Tg32 mouse model expressing hFcRn for our PK study. Mice received a single IV dose of 30 µg RNA-LNPs encoding PGT121 RibobNAb or PGT121-LS RibobNAb and blood was sampled over a 33-day period from alternating subgroups (Figure 3B).

In hFcRn-expressing mice, PGT121-LS RibobNAb demonstrated superior pharmacokinetics compared to wild-type PGT121 RibobNAb. The LS modification resulted in an approximate 2-fold extension of the serum half-life of PGT121-LS RibobNAb (PGT121 RibobNAb t_1/2_: 4.6 days vs. PGT121-LS RibobNAb t_1/2_: 8.6 days) and 1.5-fold increased systemic exposure in terms of area under the curve (AUC). The PK enhancement was hFcRn-dependent, as evidenced by the 5.6-day half-life extension observed in the NSG Tg32 mice compared with NSG mice. In contrast, the non-LS version showed no improvement.

Taken together, these findings illustrate that the RibobNAbs were successfully expressed *in vivo* and highlight the potential of LS-modified RibobNAbs to achieve favorable PK profiles in therapeutic settings.

### PGT121-LS RibobNAb matches PGT121-LS reference protein PK and antiviral characteristics in non-human primates (NHPs)

To translate the findings obtained in NSG Tg32 mice to an animal model more similar to humans, NHPs were used to directly compare the lead candidate, PGT121-LS RibobNAb, to the recombinantly expressed and purified PGT121-LS reference protein. The PK and tolerability of PGT121-LS RibobNAb were assessed in male rhesus macaques (n = 3 per group) following a single IV or IM injection of RNA-LNP at doses of 0.5 and 1.6 mg/kg. The PGT121-LS reference protein was administered IV at 1 mg/kg. Animals receiving the IM injection were observed for 120 days, while those receiving the IV injection were monitored for 180 days (Figure 4A). PK parameters were determined using noncompartmental analysis (NCA) (Figure 4B).

**Figure 4:**
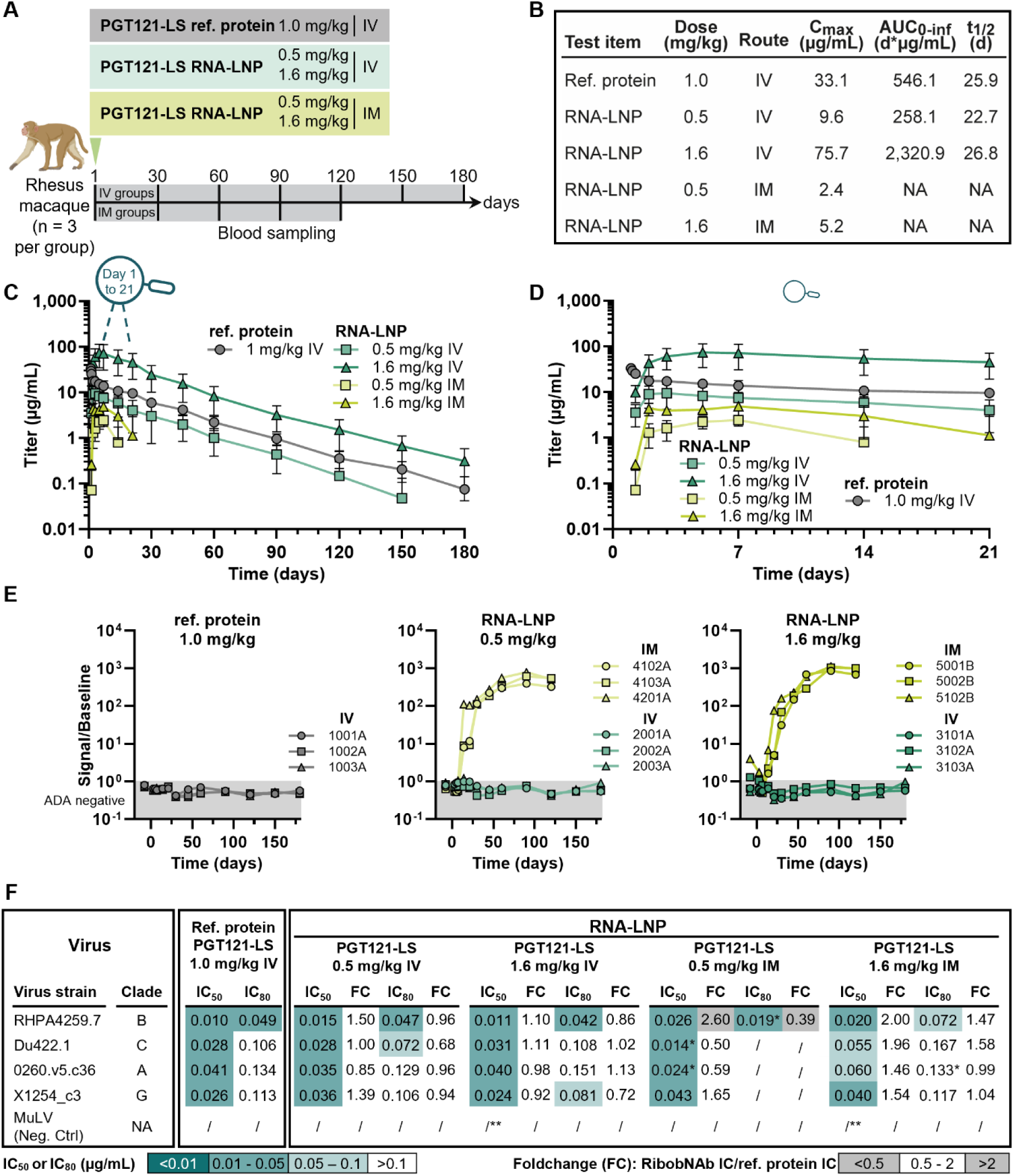
RNA-LNP-mediated PGT121-LS RibobNAb expression achieves pharmacokinetic and neutralization activity comparable to reference protein in NHPs. A) PK analysis of PGT121-LS RibobNAb compared with reference protein PGT121-LS in NHPs. The PK of PGT121-LS RibobNAb was assessed in male rhesus macaques (n = 3 per group) following a single IV or IM RNA-LNP injection at doses of 0.5 and 1.6 mg/kg. IM-treated animals were observed for 120 days and IV-treated animals were observed for 180 days. PGT121-LS reference protein was administered IV at 1 mg/kg. Serum samples were obtained at designated time points. RibobNAb titers were determined using BAMA. Error bars indicate the SD of the mean of three animals. B) PK parameters determined by NCA. C) PGT121-LS RibobNAb and reference protein titers in NHP serum over the study duration (120 days for IM-treated animals and 180 days for IV-treated animals). Detailed view of the first 21 days of the study is shown in D. D) Early PK profile (Days 0-21). Expanded view of serum PGT121-LS RibobNAb and reference protein titers during the first 21 days of the study (full study duration shown in C). E) Biotin/sulfo-tag-based bridging ADA screening assay of serum samples. Samples above signal/baseline threshold (dotted line) are considered ADA-positive. Data for individual animals are shown. F) TZM-bl neutralizing antibody assay with NHP serum samples taken on Day 2 of the study. Serum samples were tested against the indicated HIV-1 Env pseudoviruses. Mean IC_50_ and IC_80_ values of three animals are shown. *IC_50_/IC_80_ values could only be determined for one or two animals. **Single outliers were removed. AUC_0-inf_ = Area under the serum concentration-time curve from time zero to infinity; C_max_ = maximum serum concentration; FC = foldchange; IC_50_ = half maximal inhibitory concentration; IC_80_ = 80% inhibitory concentration; IM = intramuscular; IV = intravenous; LNP = lipid nanoparticle; ref. = reference; neg. ctrl. = negative control; RibobNAb = RNA-encoded, LNP-formulated anti-HIV-1 broadly neutralizing antibody; RNA = ribonucleic acid; t_1/2_ = elimination half-life.

Intravenous RNA-LNP administration resulted in dose-dependent RibobNAb expression (Figure 4C). PGT121-LS RibobNAb was quantifiable at the first sampling time point (6 h post-dose), with peak serum concentrations achieved between 24 h and 7 days post-injection (0.5 mg/kg, mean C_max_: 9.6 µg/mL, mean t_max_: 48 h; 1.6 mg/kg, mean C_max_: 75.7 µg/mL, mean t_max_: 96 h) (Figure 4B, C, D). The increase in serum concentrations observed through Days 3–7 is consistent with the kinetics of LNP-mediated RNA delivery, including LNP distribution, cellular uptake, RNA endosomal escape and translation and antibody secretion. ^6–8^ Once in the systemic circulation, IV-administered PGT121-LS RibobNAb exhibited a PK profile comparable to that of the PGT121-LS reference protein (Figure 4C, D).

RibobNAb levels were detectable for 150 days in the IV low-dose groups and throughout the entire 180-day study period in the IV high-dose group. Notably, RibobNAb half-lives (22–27 days) were comparable to that of the reference protein (25.9 days), indicating similar antibody stability and clearance kinetics (Figure 4B). As previously reported for RNA-encoded proteins, a greater-than-dose-proportional increase in exposure was observed, with a 9-fold increase in AUC following IV administration of RNA-LNPs at 1.6 mg/kg compared with 0.5 mg/kg.^32^

Translational efficiency assessment indicated that a 0.8 mg/kg dose of RNA-LNP yielded comparable systemic exposure as a 1 mg/kg dose of reference PGT121-LS.

Similarly, IM administration led to successful expression of PGT121-LS RibobNAb, detectable 6 h post-dose, at both RNA-LNP dose levels tested (Figure 4C, D). As expected, peak concentrations in the systemic circulation were delayed compared with IV administration, reaching 4– to 15-fold lower C_max_ values on Day 7 (0.5 mg/kg, mean C_max_: 2.4 µg/mL, mean t_max_: 144 h; 1.6 mg/kg, mean C_max_: 5.2 µg/mL, mean t_max_: 144 h) (Figure 4B). After reaching peak serum concentrations, RibobNAb titers decreased rapidly, correlating with ADA development in both IM groups (Figure 4E, F). As a result, estimation of PK parameters was limited to C_max_.

ADA presence and titers were determined using the previously described three-tiered ADA characterization assay.^33^ ADA were detected in all IM-treated animals from Day 14 and persisted through Day 120, with increasing response over time (Figure 4E).

Quantitative analysis revealed peak ADA concentrations at the end of the observation period (Day 120) (Supplementary Figure 1). In comparison, no ADA signals were detected in any of the IV-treated animals, regardless of the test item, suggesting that the administration route significantly influences immunogenicity. These findings demonstrate that, for the tested LNP formulation, IV is the most suitable route of administration, which is consistent with its design and optimization for IV use.

To confirm the functional activity of *in vivo*-produced PGT121-LS RibobNAb, serum samples collected on Day 2 were assessed for HIV-1 neutralization activity against a panel of strains representing multiple clades (A, B, C, and G) (Figure 4G). PGT121-LS RibobNAb demonstrated potent neutralization of the tested strains, with IC_50_ and IC_80_ values comparable to those of PGT121-LS reference protein independently of dose or administration route. Fold-change (FC) analysis revealed that PGT121-LS RibobNAb maintained equivalent potency to the reference protein for most virus strains (FC: 0.5–2.0), with all observed differences falling within the assay variation limits. Collectively, the data show the potential of RNA-LNP to deliver PGT121-LS RibobNAb, with neutralization potency and PK characteristics identical to directly administered PGT121-LS protein.

### PGT121-LS RibobNAb is well tolerated in NHPs

In addition to evaluating PK and functionality, the safety profile of PGT121-LS RibobNAb was assessed for IV and IM administration routes. Of note, the RNA-LNP formulation used in this study was designed and tested for IV delivery. Overall, PGT121-LS RibobNAb was clinically well tolerated regardless of the administration route, with no mortalities and only transient treatment-related effects (Figure 5A).

**Figure 5:**
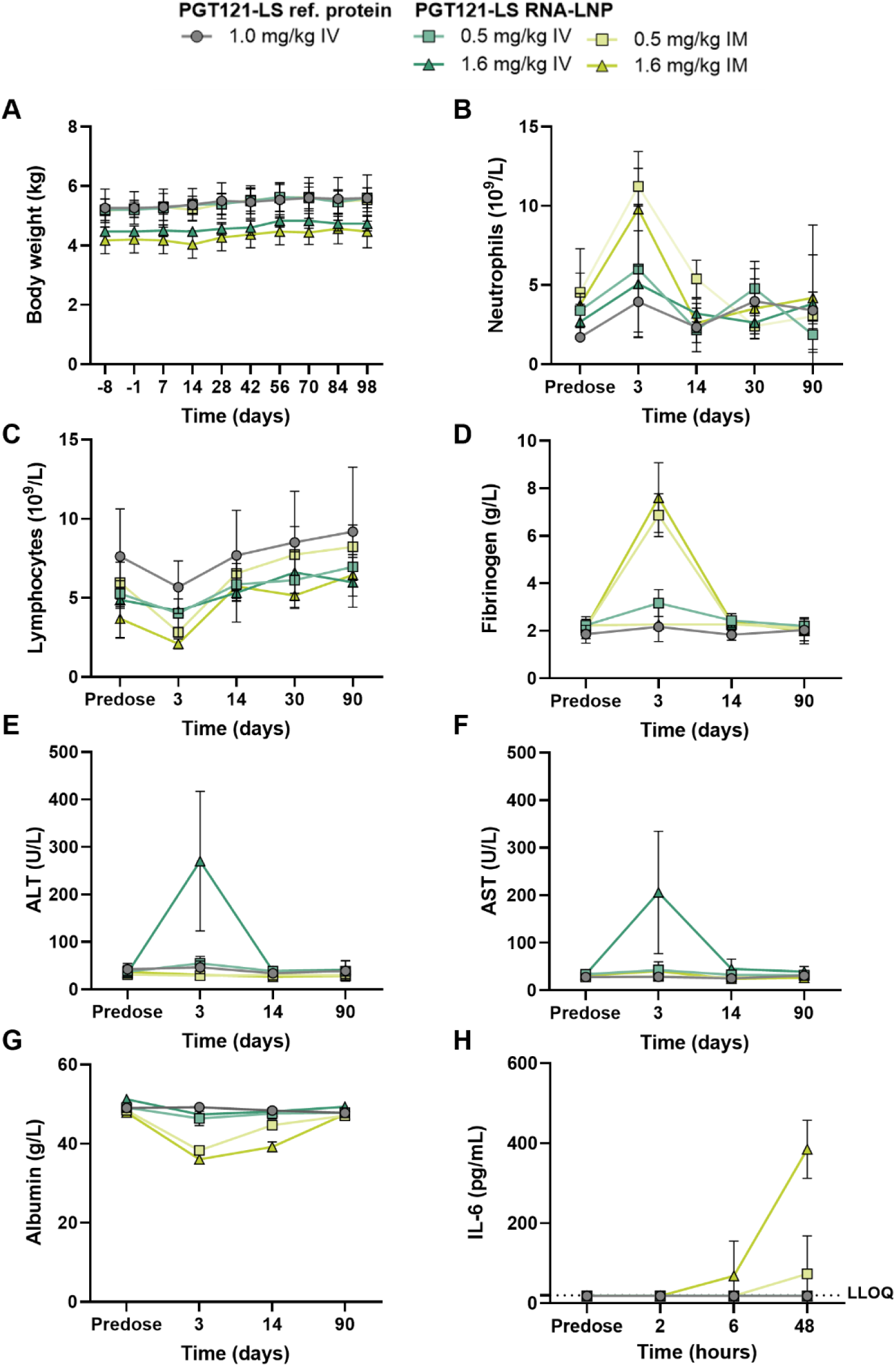
PGT121-LS RibobNAb is well tolerated by NHPs. A) Body weights of male rhesus macaques (n = 3 per group) following a single IV or IM injection of PGT121-LS RNA-LNP injection at doses of 0.5 and 1.6 mg/kg, or a single IV injection of PGT121-LS reference protein at a dose of 1.0 mg/kg. The body weights were measured for all animals prior to group assignment and at the indicated time points throughout the study. B) Neutrophil counts, C) lymphocyte counts and D) fibrinogen levels in peripheral blood from NHPs that received the treatments at predose and on Day 3, 14 and 90 post-injection (B-D). E) ALT, F) AST and G) albumin levels in serum of NHPs that received the treatments at predose and on Day 3, 14 and 90 post treatment (E-G). H) IL-6 serum concentrations were quantified predose and 2, 6 and 48 h post-administration in peripheral blood of NHPs via multiplex assay. A-H) Error bars indicate the SD of the mean of three animals. ALT = alanine aminotransferase; AST = aspartate aminotransferase; IL-6 = interleukin-6; IM = intramuscular; IV = intravenous; LNP = lipid nanoparticle; LLOQ = lower limit of quantification; ref. = reference; RibobNAb = RNA-encoded, LNP-formulated anti-HIV-1 broadly neutralizing antibody; RNA = ribonucleic acid.

However, IM administration was associated with a localized inflammatory reaction at the injection site, and with pyrexia in all animals receiving the high dose (1.6 mg/kg), which resolved within one week (Figure S2A). IM administration also induced hematological changes associated with acute inflammation, including a transient, non-dose-dependent increase in neutrophil counts (1.5– to 6-fold) on Day 3 which returned to baseline by Day 14 (Figure 5B), along with a slight decrease in lymphocyte counts that remained within the historical reference range of the testing site (Figure 5C). Other hematological changes were deemed incidental, as they either fell within the range of pre-treatment values or reflected normal inter-animal variability for this species (Table S1).

IM dosing caused a 2.5– to 4-fold, non-dose-dependent increase in the acute-phase protein fibrinogen on Day 3 compared with pre-treatment values, indicating an inflammatory response. This increase was transient, with fibrinogen levels returning to baseline by Day 14 in all animals (Figure 5D).

While liver enzymes remained at baseline levels following IM administration (Figure 5E, F), a dose-dependent decrease in serum albumin and a slight increase in serum globulin were observed, resulting in a lower A/G ratio indicative of inflammation, which showed a trend towards recovery by Day 14 (Figure 5G and Figure S2B). Other observed minor changes in clinical chemistry values throughout the study were considered incidental or reflected normal inter-animal variation in this species (Table S2 to Table S3).

In addition, IM administration at 1.6 mg/kg was associated with an increase in the inflammatory cytokine interleukin-6 (IL-6) at 48 h post-dose correlating with fever and localized inflammation, while other cytokines showed sporadic or negligible changes unrelated to PGT121-LS RNA-LNP treatment (Figure 5H).

In contrast, IV administration was not associated with local reactogenicity, fever, or acute-phase protein changes (Figure 5B to D). IV dosing was characterized by transient, dose-dependent increases in liver enzymes, with alanine aminotransferase (ALT) levels rising 3– to 14-fold (Figure 5E) and aspartate aminotransferase (AST) levels increasing 3– to 10-fold (Figure 5F) from baseline following administration of the higher dose (1.6 mg/kg). These elevations peaked on Day 3 and returned to baseline by Day 14, suggesting low-grade, reversible liver stress. Transient, self-limiting elevations in liver enzymes have been reported previously with RNA-LNP approaches.^34–36^ Notably, IV administration of RNA-LNP, such as the formulation used in our study, is designed to target the liver, resulting in efficient protein production.^6–8^

Complement activation was more pronounced with IV administration, with dose-dependent increases in plasma Bb and sC5b-9 complement levels observed at 30 minutes (min) and 2 h post-dose, returning to baseline by 24 h. IM administration induced only minor complement activation at 24 h in high-dose animals. Fold changes of Bb and sC5b-9 complement are summarized in Table 1 and Table 2.

**Table 1:** Summary of fold changes of Bb complement.

| Time points | Reference IV |  | 0.5 mg/kg IV |  | 1.6 mg/kg IV |  | 0.5 mg/kg IM |  | 1.6 mg/kg IV |  |
| --- | --- | --- | --- | --- | --- | --- | --- | --- | --- | --- |
|  | Min | Max | Min | Max | Min | Max | Min | Max | Min | Max |
| Pre-treatment | - | - | - | - | - | - | - | - | - | - |
| 30 min | - | - | 1.6 | 2.6 | 1.9 | 4.9 | - | - | - | - |
| 2 h | - | - | 1.5 | 3.1 | 2.6 | 3.6 | - | - | - | - |
| 24 h | 1.1 | 1.1 | 1.1 | 1.2 | 1.3 | 2.6 | - | - | 1.2 | 1.5 |

Min = minimum; Max = maximum; IM = intramuscular; IV = intravenous.

**Table 2:** Summary of fold changes of sC5b-9 complement.

| Time points | Reference IV |  | 0.5 mg/kg IV |  | 1.6 mg/kg IV |  | 0.5 mg/kg IM |  | 1.6 mg/kg IV |  |
| --- | --- | --- | --- | --- | --- | --- | --- | --- | --- | --- |
|  | Min | Max | Min | Max | Min | Max | Min | Max | Min | Max |
| Pre-treatment | - | - | - | - | 1.1 | 1.1 | - | - | - | - |
| 30 min | - | - | 1.5 | 2.0 | 4.7 | 7.2 | - | - | - | - |
| 2 h | - | - | 1.5 | 3.3 | 4.0 | 12.5 | - | - | - | - |
| 24 h | - | - | 1.2 | 1.3 | 2.1 | 8.2 | 1.1 | 1.6 | 1.5 | 2.3 |

Min = minimum; Max = maximum; IM = intramuscular; IV = intravenous.

Overall, IV administration demonstrated a more favorable safety profile compared with IM administration, with no local or systemic responses. While transient liver enzyme elevations and complement activation were observed with IV delivery, these changes were reversible and resolved within two weeks. These findings suggest that, for the formulation used in this study, IV administration is the optimal route for systemic delivery of PGT121-LS RibobNAb.

### Population PK modeling and human dose prediction

Building on the favorable PK profile of PGT121-LS RibobNAb in NHPs, we designed a population PK model for FIH dose prediction. We developed a model capturing the distinct kinetics of RNA-LNP-mediated protein expression (Figure 6A). Whereas antibodies delivered as proteins are directly bioavailable upon administration followed by tissue distribution and elimination, RNA-LNP administration is followed by LNP distribution, cellular uptake, RNA endosomal escape and RNA translation, resulting in delayed but transiently sustained RibobNAb production from host cells, e.g. liver cells, with peak concentrations reached several hours to days post-administration.

**Figure 6:**
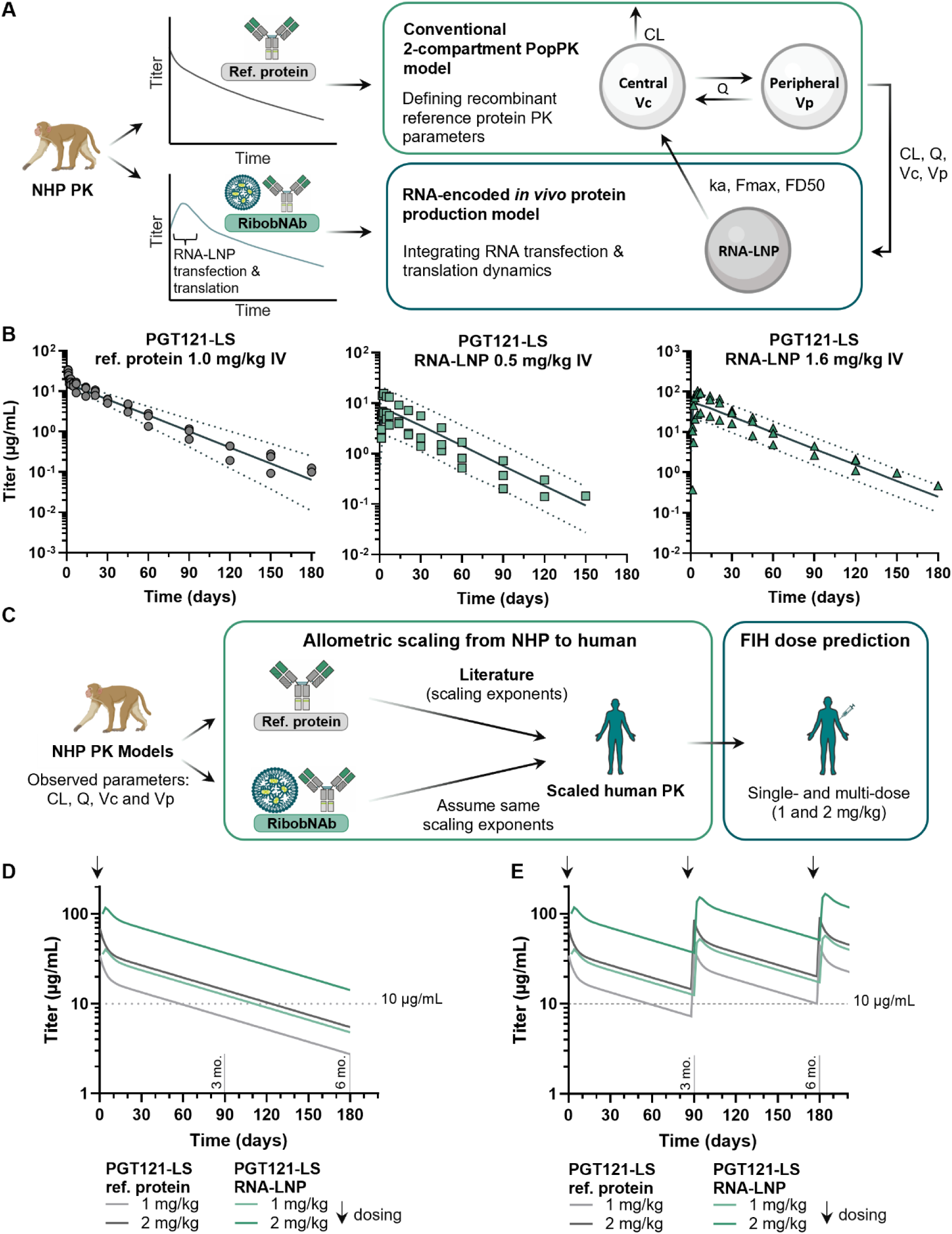
PopPK modeling and dose prediction indicates that therapeutic titers with long-term exposure can be achieved in humans. A) Modeling workflow: Antibodies administered as purified proteins and RibobNAbs show distinct PK profiles in NHPs. Purified protein antibodies are directly bioavailable upon administration followed by immediate elimination. RNA-LNPs require cellular uptake and translation, resulting in delayed RibobNAb production with peak concentrations reached several hours to days post-administration. NHP pharmacokinetic data informed development of conventional two-compartment PopPK models for reference protein and RNA-encoded *in vivo* protein production models incorporating translation dynamics (k_a_, Fmax, FD50) for RibobNAbs. B) Observed NHP pharmacokinetic profiles and model predictions for PGT121-LS reference protein (1.0 mg/kg IV) and PGT121-LS RibobNAb encoded by RNA-LNP formulations (0.5 and 1.6 mg/kg IV). Symbols represent individual animals; solid lines show population predictions with 90% prediction intervals (dotted lines). C) Allometric scaling approach: NHP-derived parameters (CL, Q, Vc, Vp) were scaled to humans using published exponents for LS-engineered antibodies, assuming equivalent scaling for RibobNAb, enabling FIH dose predictions. D-E) Human dose predictions targeting 10 μg/mL therapeutic threshold (dashed line): single-dose (D) and monthly dosing regimens (E) for reference protein and RNA-LNP at 1 and 2 mg/kg. Simulations demonstrate sustained therapeutic coverage with both platforms but better coverage with the RibobNAb. CL = clearance (elimination clearance); FD50 = dose producing 50% of maximal translation; Fmax = maximum translation; FIH = First-in-Human; IV = intravenous; k_a_ = translation rate; LNP = lipid nanoparticle; NHP = non-human primate; PK = pharmacokinetics; ref. = reference; PopPK = population PK; Q = intercompartmental clearance (distribution clearance between compartments); RibobNAb = RNA-encoded, LNP-formulated anti-HIV-1 broadly neutralizing antibody; RNA = ribonucleic acid; t_1/2_ = elimination half-life; Vc = central volume of distribution; Vp = peripheral volume of distribution.

Accordingly, we first fitted a conventional two-compartment population pharmacokinetics (PopPK) model from our reference protein dataset. We defined structural parameters including clearance (CL), intercompartmental clearance (Q), central volume (Vc), and peripheral volume (Vp). These parameters were incorporated into an RNA-LNP-specific model accounting for RNA-LNP delivery, cellular transfection, RNA translation dynamics, and subsequent antibody production, described as translation rate constant (k_a_) and dose-dependent translational efficiency (F). The fully parameterized model integrates sequentially the RNA-LNP kinetic model followed by a two-compartment model resolving the antibody PK.

With the model structure established, we proceeded to evaluate its fit and estimate RNA-LNP-specific parameters. Visual predictive checks (VPCs) for dose-stratified models confirmed adequate model fit (Figure 6B), while inter-individual variability was identified on CL, Vc and Vp. For RNA-encoded antibody PK modeling, structural parameters (Table S4) were fixed and only k_a_ and F were estimated. Non-linear translational efficiency ranging from 0.8 to 2.4 (for low and high doses, respectively) was observed. The parameter estimates for the model are shown in Table S4.

Human dose projections for the RNA-LNP were generated by applying established allometric exponents for LS-engineered IgG to the previously characterized structural pharmacokinetic parameters (CL, Q, V_c_, V_p_) while assuming comparable translation-related parameters (k_a_ and F) between NHP and humans.

Our model shows that a single 2 mg/kg dose of RNA-LNP could sustain a titer >10 µg/mL for over 6 months, a value described as therapeutic threshold in the clinic.^37^ Similarly, a 2 mg/kg IV dose of PGT121-LS reference protein is expected to maintain therapeutic concentrations for about 4 months (Figure 6D). Simulations using repeated dosing with a 3-month interval predict antibody accumulation that would allow dose reduction while maintaining titers above therapeutic concentrations (Figure 6E).

## Discussion

As a promising alternative to cART, HIV bNAbs combine viremia suppression with the added potential of engaging host immunity for possible durable viral control.^38–40^ However, high costs and scalability limitations associated with antibody manufacturing hinder their broad deployment, driving interest in alternative delivery platforms.^4,41^

The utility of vector-mediated antibody delivery has been demonstrated using adenoviral vectors and DNA-based systems, which have achieved *in vivo* antibody production for HIV and other indications.^42–45^ More recently, advances in RNA technology have positioned RNA-encoded antibodies as an especially attractive platform, enabling transient, controllable protein expression without genomic integration and with the potential to reduce immunogenicity associated with the repeat administration of viral vectors.^46–49^ RNA-based antibody delivery has already shown promise across cancer immunotherapy, neurodegenerative diseases, and infectious diseases.^50–52^ Phase I clinical-stage examples include BNT142 (NCT05262530), an RNA-encoded bispecific antibody for the treatment of solid tumors, and mRNA-1944 (NCT03829384), which encodes a neutralizing antibody against chikungunya virus, demonstrating that RNA-LNP formulations can safely and effectively drive *in vivo* expression of diverse antibody constructs in humans.^50,52^

In this study, we used *in vitro*-transcribed RNA as a flexible platform for testing Fc-engineered versions of the potent, clinically-tested HIV bNAb PGT121. The Fc-modified variants incorporated PK-enhancing mutations designed to prolong half-life, with the aim of reducing the frequency of administrations required to maintain viral suppression in PWH. We successfully expressed four PGT121 variants *in vitro* that retained sub-nanomolar binding affinity to SOSIP and demonstrated viral neutralization comparable to PGT121 reference protein, supporting the use of RNA to streamline the production and screening of functional bNAbs. While we focused on Fc modifications on IgG1 in this work, our pipeline is also amenable to isotype switching as demonstrated by Rouzeau et al.^12^

Single-dose IV administrations of PGT121-LS RNA-LNP (0.5 and 1.6 mg/kg) were well tolerated in male rhesus macaques, with only transient and reversible AST/ALT elevations. The PK and antiviral activity of IV-delivered PGT121-LS RibobNAb were indistinguishable from PGT121-LS reference protein. These observations are consistent with previous work by Bähr-Mahmud et al., who reported no toxicity in rodents and no safety concerns in NHPs when comparing the protein anti-claudin 18.2 antibody IMAB362 with RNA-LNP-encoded RiboMab01.^6^ In the HIV field, others have successfully expressed RNA-encoded anti-HIV bNAbs in various organisms, including mice and sheep.^53,54^ Importantly, in our single-dose IV setting, we did not detect ADA elicitation over the course of 180 days. However, Rosenberg et al. observed ADA formation in infant macaques after repeated subcutaneous injection of PGT121-LS protein.^55^ Whether multiple IV injections of PGT121-LS RNA-LNP lead to ADA elicitation still needs to be investigated.

While our LNP delivery system was optimized for low immunogenicity and IV administration for efficient protein production in the liver, IM administration offers practical advantages in areas with limited access to trained healthcare personnel and infusion equipment.^56,57^ Thus, we sought to assess the potential of our formulation for IM delivery and conducted a side-by-side comparison of IV and IM delivery in NHPs. Compared to IV administration, IM delivery demonstrated reduced bioavailability and elevated immunogenicity. Several mechanistic factors may contribute to this suboptimal performance. The prolonged absorption required for LNPs to enter systemic circulation and reach the liver compromises efficacy, as the liver is the primary production organ for our LNP formulation clinically validated for IV delivery.^6–8^ The observed delay in peak serum concentration could reflect LNP retention at the injection site with prolonged release into systemic circulation. Alternatively, this observation could be due to delayed release of RibobNAb produced in muscle cells, similar to what is observed after subcutaneous injection of an antibody.^58,59^ Notably, the fractional synthetic rate of muscle cells is 10– to 50-fold lower than that of hepatocytes (0.02–0.09%/h vs 0.97–0.99%/h).^60,61^ The observed 3– to 10-fold reduction in C_max_ post-IM dosing could be attributable to lower transfection efficiency and/or reduced protein production.

In addition, the rapid development of ADAs in IM-dosed animals likely contributed to accelerated RibobNAb clearance. The induction of ADA coincided with an elevation in pro-inflammatory markers and modulation of acute-phase proteins, indicating an immune response likely promoting their elicitation. Overall, the reduced performance associated with IM administration may stem from formulation-specific limitations rather than platform constraints or PGT121-LS immunogenicity, suggesting that optimizing the LNP composition could improve systemic bioavailability after intramuscular injection.

To evaluate long-term efficacy, we developed a PK model tailored to the unique features of RNA-LNP-encoded antibodies, capturing LNP distribution, cell transfection, RNA translation, RibobNAb secretion, and subsequent RibobNAb PK. Our empirically driven framework was designed to focus on systemic exposure with reduced structural and data complexity, supporting practical dose projections and clinical translation. Despite differences in scope and granularity, our conclusions align with a recently published mechanistic model that further characterizes intracellular trafficking, translation, and protein turnover, in oncology settings.^62^ In addition, our prediction for the PGT121-LS reference protein corresponds well with clinical PK data obtained for another recombinant protein variant of PGT121 (PGT121.414.LS).^63^

Our human PK model predicts that maintaining therapeutic serum levels of RibobNAb above 10 µg/mL would require RNA-LNP doses of 2 mg/kg and 1 mg/kg every 6 and 3 months, respectively If translational efficiency were reduced in humans (e.g., allometric exponent of 0.7 for k_a_ and F), a dose of 5 mg/kg would be required, which may not be clinically feasible because of the toxicity limits of currently validated LNP formulations (data not shown). As our study used simian-human immunodeficiency virus (SHIV)-uninfected NHPs, the pharmacokinetics of PGT121-LS RibobNAb may differ in PWH, who often experience chronic inflammation even when virus is suppressed by cART.^64–66^ Notably, previous clinical data showed a faster clearance of recombinantly expressed and purified PGT121 in PWH compared with PWoH,^17^ supporting the need for dedicated PK and efficacy studies of RibobNAbs in an *in vivo* SHIV/HIV infection model.

Besides maintaining therapeutic serum levels, RNA-encoded antibody strategies must account for the emergence of resistance to single bNAbs.^17,67–70^ bNAb-based therapy therefore requires the administration of multiple complementary bNAbs to significantly enhance viral suppression efficacy and delay viral rebound compared to monotherapy.^18,19,37,70^ RNA-LNP platforms are particularly well-suited for implementing these combination strategies, as they can encode and co-deliver multiple bNAbs.^11,71^

In summary, our work demonstrates that RNA-encoded PGT121-LS RibobNAb can achieve therapeutically relevant exposure with favorable PK and low immunogenicity in NHPs upon IV administration. PK modeling and allometric scaling support the potential for clinically meaningful dosing intervals in humans.

## Material and methods

### Study design

This study describes the preclinical pharmacology of PGT121 RibobNAb variants, both without (PGT121 RibobNAb) or with Fc modifications (PGT121-LS, PGT121-del294, PGT121-LS-del294 RibobNAb). PGT121 RibobNAbs are LNP-formulated, RNA-encoded HIV-1 bNAbs targeting a V3-glycan-dependent site on gp120. *In vitro*, cellular uptake of RNA and RNA-LNPs encoding PGT121 RibobNAbs, and subsequent translation, assembly, and secretion were assessed in cell culture experiments by ELISA and Western blot. Binding to SOSIP was evaluated using BLI and ELISA, and neutralizing activity of *in vitro*-expressed RibobNAb variants was assessed in a pseudovirus neutralization test (pVNT). *In vivo,* expression of PGT121 RibobNAb variants was assessed in NSG mice. The PK of the lead candidate, PGT121-LS RibobNAb, was further evaluated in NSG Tg32 mice and rhesus macaques. Tolerability of PGT121-LS RibobNAb in NHPs was investigated by clinical pathology assessments, and the functional activity of *in vivo*-expressed PGT121-LS RibobNAb was further assessed using a pVNT.

For the mouse experiments, animals were randomly assigned to study groups. Investigators were not blinded to group assignments during experimentation and analysis. Federation of European Laboratory Animal Science Associations (FELASA), Gesellschaft für Versuchstierkunde/Society for Laboratory Animal Science (GV-SOLAS), and local regulatory guidelines were adhered to in the planning, conduct, and reporting of mouse experiments. Specific details regarding the number of animals per group and route of administration are provided in figure legends, as applicable.

For the NHP experiments, animals were initially assigned to dose groups during the acclimation period using a randomized stratification system based on body weight. Subsequently, a reassignment of animals was conducted based on dose levels. Animals expected to reach a body weight of 4.8 kg or higher at the start of dosing were excluded from the high-dose group.

### Animals and husbandry

Female and male NSG mice (NSG; Janvier Labs, Le Genest-Saint-Isle, France; RRID:MGI:6357755) as well as NSG hFcRn Tg32 mice (NSG hFcRn Tg32; Charles River Labs, Wilmington, Massachusetts; RRID:IMSR_JAX:005557) were at least 6 weeks old at the start of the study and had an average body weight ∼25 g. The mice were housed in individually ventilated cages with weekly bedding changes and were acclimatized for one week prior to the start of experiments.

All mouse studies were approved by the regional animal welfare committee (ethical approval no. G 17-12-001) and conducted in accordance with FELASA recommendations, the German Animal Welfare Act, and EU Directive 2010/63/EU. Measures were taken to minimize pain and suffering of the animals throughout the study. Additionally, euthanasia was performed in adherence to the guidelines of the GV-SOLAS, using cervical dislocation or CO_2_ asphyxiation at the end of an experiment or when humane endpoints were reached.

The NHP study was conducted at Charles River Laboratories Montreal ULC in Laval, Canada. Sixteen non-naïve male rhesus monkeys, including one spare animal (originally received from ISAGE, Kunming, China) were transferred to the study site at least 12 days prior to the start of the study. At the onset of dosing, the age of the animals ranged from 4 to 6 years, and the body weights ranged from 3.7 to 5.9 kg. Prior to transfer from the colony, all animals were subjected to a health assessment and tested at least once for tuberculosis by intradermal injection of tuberculin. An anthelmintic treatment was administered to each animal by subcutaneous injection. The evaluations were performed in accordance with the standard operating procedures by technical staff. The results of the evaluations were reviewed by the Clinical Veterinarian to ensure that animals were healthy and suitable for use in the study. Animals were housed in groups of three animals of the same treatment group in stainless steel monkey cages equipped with an automatic watering system. The room temperature was maintained at 21 °C ± 3 °C (maximum range) and the relative humidity at 50% ± 20%. Rooms were alternately lit and darkened in a 12-hour dark/12-hour light cycle. A standard certified commercial chow (Envigo Teklad Certified Hi-Fiber Primate Diet #7195C) was provided to the animals twice daily. Treats (fruits and/or vegetables) were provided to the animals as part of the facility enrichment program. Drinking water was offered *ad libitum*.

#### In life observations in NHPs

Mortality checks were conducted at least twice daily throughout all phases of the study. Body weights were measured for all animals at least once prior to assignment, weekly from Day –8 to Day 15 (second week post-dose), and bi-weekly thereafter. Measurements continued until Day 119 for IM groups, and until Day 169 for IV groups.

Clinical pathology evaluations, including hematology, clinical chemistry, and coagulation, were performed once during the pre-dose period and on Days 3, 14, and 90 for all animals. Hematology assessments were also conducted on Day 30. Blood samples were collected via femoral venipuncture, with food removed overnight prior to collections for clinical chemistry analyses.

For complement analysis, blood samples were collected from all animals during the pre-treatment period and at 30 min, 2 h, and 24 h post-dose on Day 1. Samples were collected via femoral venipuncture into K2 EDTA tubes and kept on wet ice prior to centrifugation (+4 °C, 1,500 × g, 10 min). Plasma was aliquoted into four separate tubes:

Aliquots 1 and 2: 40 μL each for Bb complement marker analysis

Aliquots 3 and 4: 80 μL each (72 μL plasma + 8 μL nafamostat mesilate [Futhan]) for sC5b-9 complement marker analysis

A Futhan complement stabilizing reagent (500 μg/mL) was pre-added to aliquots 3 and 4 for a final concentration of 50 μg/mL. Samples were stored at –70 °C until analysis. Complement markers Bb and sC5b-9 were analyzed using validated ELISA assays at Charles River Laboratories Montreal ULC. Validation was performed in cynomolgus monkey plasma, and feasibility in rhesus monkey plasma was confirmed through *in vitro* stimulation with Zymosan. Plasma from untreated rhesus monkeys served as positive and negative controls for parallelism assessments.

For cytokine analysis, blood samples were collected from all animals during the pre-treatment period and at 2, 6, and 48 h post-dose. Samples were collected via femoral venipuncture into K2 EDTA tubes and kept on wet ice prior to centrifugation (+4 °C, 1,500 × g, 10 min). Plasma was aliquoted into two tubes, stored at –70 °C until analysis.

Cytokine analysis was performed using the following assays:

TNF-α, IFN-γ, IL-1β, IL-2, IL-6, IL-8, and IL-10, using a multiplex (Millipore kit, catalogue No. PRCYTOMAG-40K-07); IFN-α using an ELISA assay (catalogue No. 41110-1, PBL assay science); and IL-12p70 using R&D kit (catalogue No. FCSTM21-02).

### Cell lines

The HEK-293T-17 cell line (ATCC no. 11268, LGC Limited) utilized for RibobNAb production via RNA transfection or RNA-LNP transfection was cultured in Dulbecco’s Modified Eagle Medium (DMEM) supplemented with GlutaMAX (Thermo Fisher Scientific, Waltham, Massachusetts) and 10% fetal bovine serum (FBS) (Thermo Fisher Scientific, Waltham, Massachusetts). Cells were maintained at 37 °C in a humidified incubator with 7.5% CO_2_.

HEK-293T-17 cells (ATCC) used for pseudovirus generation were maintained at 37 °C and 5% CO_2_ in DMEM (Thermo Fisher Scientific, Waltham, Massachusetts) supplemented with 10% FBS (Sigma-Aldrich, St. Louis, Missouri), 1 mM sodium pyruvate, 2 mM L-glutamine, and 1x antibiotic-antimycotic (all from Thermo Fisher Scientific, Waltham, Massachusetts).

TZM-bl cells (NIH AIDS Reagent Program) used in pVNT assays were maintained at 37 °C in 5% CO_2_ in DMEM (Thermo Fisher Scientific, Waltham, Massachusetts) supplemented with 10% FBS (Sigma-Aldrich, St. Louis, Missouri), 1 mM sodium pyruvate (Thermo Fisher Scientific, Waltham, Massachusetts), 2 mM L-glutamine (Sigma-Aldrich, St. Louis, Missouri), 50 µg/mL gentamicin (Sigma-Aldrich), and 25 mM 4-(2-hydroxyethyl)-1-piperazineethane-sulfonic acid (HEPES, Thermo Fisher Scientific, Waltham, Massachusetts or Sigma-Aldrich, St. Louis, Missouri).

### Generation of PGT121 RNA and RNA-LNP variants

PGT121-encoding RNAs were generated as previously described^8^ via the construction of human codon-optimized DNA sequences as well as *in vitro* transcription of DNA sequences to generate the individual heavy and light chain-encoding RNAs. The heavy and light chain variable domain sequences of PGT121 as derived from the literature^15^ were subcloned into the multiple-cloning site (MCS) of pST1-hAg-Kozak-MCS-F-I-A30LA70 (modified from^72^), containing the respective human IgG1 constant heavy and light chain domain sequences encoding PGT121 RibobNAb variants both without or with the indicated Fc modifications. In addition, the RNA transcripts incorporate a set of conserved structural features that promote RNA stability and efficient protein translation. These regulatory elements consist of a CC413 cap at the 5’ end, a 5’-untranslated region (AGA-dEarI-hAg), a 3’-untranslated region containing the FI element, and a 100-nucleotide poly-adenine tail with an integrated linker at the 30th position (A30L70).

RibobNAb-encoding DNA sequences were transcribed *in vitro* using T7 RNA polymerase (Ambion from Thermo Fisher Scientific, Waltham, Massachusetts), modified nucleotide triphosphates, and CleanCap 413 (TriLink Biotechnologies, San Diego, California) for 5’ capping, producing N1-methylpseudouridine (m1Ψ)-capped RNA. The RNA products were isolated using magnetic bead separation, followed by cellulose purification, resuspension in ribonuclease (RNase)–free water (B. Braun, Melsungen, Germany) containing 10 mM HEPES/0.1 mM EDTA (BioChrom), and storage in nuclease-free reaction tubes (Eppendorf, Hamburg, Germany). Quality control of the synthesized RNAs was performed using capillary electrophoresis (Agilent 2100 Bioanalyzer, Agilent Technologies, Santa Clara California) and concentration determined by spectrophotometry (NanoDrop 2000c, Thermo Fisher Scientific, Waltham, Massachusetts). PGT121 RNAs were then stored at −65 to −85 °C and either used for RibobNAb expression via RiboJuice™ or for formulation into LNPs. To generate PGT121 RNA-LNP variants, the respective heavy– and light-chain-encoding PGT121 RNAs were mixed at a ratio of 1.5:1 and encapsulated in an LNP formulation consisting of an ionizable cationic lipid, a polyethylene glycol (PEG) lipid, phospholipid, and cholesterol, followed by storage at −65° to −85 °C. The formulations were characterized with regard to particle size, encapsulation efficiency, and RNA concentration.

### RiboJuice™ transfection

HEK-293T-17 cells were thawed and passaged at least three times prior to use. Cells were seeded at a density of 4.0 × 10^5^ cells/mL/well in a cell culture 12-well plate (Greiner Bio-One, Kremsmünster, Austria) 24 h before the transfection and incubated overnight at 37 °C with 7.5% CO_2_. To express the RibobNAbs, 1 µg RNA were transfected for each well using a heavy chain to light chain ratio of 1.5:1 (0.6 µg heavy chain /0.4 µg light chain). RiboJuice™ RNA transfection reagents were prepared according to manufacturer’s instructions (Merck Millipore, Billerica, Massachusetts).

Complexes with 1 µg RNA were dropwise added to each well and afterwards mixed by gently shaking the plate. The cells were incubated at 37 °C in a humidified incubator with 7.5% CO_2_ for 48 h. At the end of the incubation period, the cell culture supernatants were centrifuged at 400 × g for 2 min and afterwards at 3,000 × g for 10 min to remove any residual cells or debris. Cell culture supernatants were stored at 4 °C until analysis.

### RNA-LNP transfection

HEK-293T-17 cells were thawed and passaged at least three times prior to use. Cells were seeded at a density of 4.0 × 10^5^ cells/mL in a cell culture 12-well plate (Greiner Bio-One, Kremsmünster, Austria) 24 h before RNA-LNP transfection and incubated overnight at 37 °C with 7.5% CO_2_. After incubation, the cell culture supernatant was carefully aspirated and replaced with 800 µL prewarmed Opti-MEM-I reduced serum medium per well (Thermo Fisher Scientific, Waltham, Massachusetts). The RNA-LNP aliquots were thawed and thoroughly mixed by gently flipping the tube without vortexing. RNA-LNPs (1 µg/well) were prepared in 200 µL Opti-MEM-I reduced serum medium (Thermo Fisher Scientific, Waltham, Massachusetts) and dropwise added to each well. The plate was gently swirled to evenly distribute the RNA-LNP onto the cells. The cells were incubated at 37 °C in a humidified incubator with 7.5% CO_2_ for 48 h. At the end of the incubation period, the cell culture supernatants were centrifuged at 400 × g for 2 min and afterwards at 3,000 × g for 10 min to remove any residual cells or debris. Cell culture supernatants were stored at 4 °C until analysis.

### ELISA for RibobNAb quantification

For quantification of RibobNAbs in cell culture supernatant, Reagent A (Gyrolab® huIgG Kit, Gyros Protein Technologies, Uppsala, Sweden), which contains a ready-to-use biotinylated derivative of protein A, was used as capture. For quantification of RibobNAbs in mouse serum Reagent A (PK or TK Kit), which contains a ready-to-use biotinylated anti-human IgG Fc-specific antibody, was used as capture. Detection was performed using Reagent B from the same kits (Gyrolab® huIgG Kit, Gyrolab® Generic PK or TK Kit), which contains a ready-to-use Alexa^®^ Fluor 647 labeled F(ab’)2 fragment of anti-human IgG. The assays were processed in a Gyrolab^®^ Bioaffy™1000 HC CD (Generic PK Kit) or Gyrolab® Bioaffy™ 20 HC CD (Generic TK Kit). Cell culture samples were diluted 10-fold in Reagent E buffer (Gyrolab® huIgG Kit) for RiboJuice™ transfected samples and for RNA-LNP transfected samples. Serum samples were diluted 10-fold with Reagent F buffer (from Gyrolab® Generic PK or TK Kit). The CD columns were washed with Reagent C and Reagent D (Gyrolab® huIgG Kit, Gyrolab® Generic PK or TK Kit).

An eleven-point, three-fold serial dilution standard curve was generated using an IgG1 reference protein (Gyrolab® Gyros Protein Technologies, Uppsala, Sweden; LakePharma, San Carlos, California; CAVD / DUKE University, Durham, North Carolina) diluted in Reagent E buffer for cell culture supernatant and Reagent F buffer for serum samples. Dilutions were made in buffers spiked with sample-matched matrix at the final dilution factor.

All data were generated on a Gyrolab® xPand ELISA device (Gyros ProteinTechnologies AB), and results analyzed with Gyrolab Evaluator software.

### SDS-PAGE and Western blotting

Cell culture supernatant samples of HEK-293T-17 cells transfected with PGT121 RibobNAb-encoding RNA variants were prepared by mixing with 4x Laemmli buffer (Bio-Rad, Hercules, California). For reducing conditions, dithiothreitol (DTT) (Carl Roth, Karlsruhe, Germany) was added to a final concentration of 100 mM. For non-reducing conditions, an equivalent volume of water was added instead of DTT. All samples were denatured by heating at 95 °C for 5 min, followed by cooling to room temperature (RT). A total of 7 ng of protein for RiboJuice™ transfected samples and 2.23 ng of protein for LNP lipofected samples together with undiluted mock or untreated control was loaded onto precast 4-15% Criterion™ gradient SDS-PAGE gels (Bio-Rad, Hercules, California) and separated by SDS-PAGE for 40 min at 250 V at 0 °C in TGS running buffer (Bio-Rad, Hercules, California). For reference, 7 ng of a recombinantly expressed and purified PGT121 (LakePharma, San Carlos, California) was included. As protein standard, 10 µL of 1:1 premixed Precision Plus Protein™ All Blue Prestained Protein Standards and Precision Plus Protein™ Unstained Protein Standards (Bio-Rad, Hercules, California) ladder was used.

For Western blotting, proteins were transferred onto a Trans-Blot Turbo Midi 0.2 µm nitrocellulose membrane (Bio-Rad, Hercules, California) at 2.5 A for 7 min using the Trans-Blot®Turbo™ (Bio-Rad, Hercules, California) blotting system. Membranes were blocked for 1 h at RT with 5% skimmed milk dissolved in Tris-buffered saline (TBS) (Bio-Rad, Hercules, California) containing 0.5% Tween-20 (TBST 0.5%, Sigma). After blocking, membranes were washed with TBS containing 0.1% Tween-20 (TBST 0.1%) and incubated with a Peroxidase-conjugated goat anti-human-Fcγ-specific detection antibody (Jackson ImmunoResearch Europe Ltd, Ely, England) diluted 1:1,000 and a Peroxidase-conjugated goat anti-human Lambda Light Chain detection antibody (Bio-Rad, Hercules, California) diluted 1:500 in TBST 0.5% supplemented with 3% BSA Fraction V (Eurobio Scientific, Paris, France). The membrane was incubated for 1 h and washed with TBS containing 0.1% Tween-20 (TBST 0.1%).

Protein bands were visualized with Clarity™ Western ECL Reagent (Bio-Rad, Hercules, California) on a Vilber Fusion FX imaging device (Vilber) with an exposure time of 0.1 s and merged with white light image to visualize marker. Data analysis was performed with the Image Lab Software (Bio-Rad, Hercules, California).

### Bio-layer interferometry

BLI measurements were performed using the Octet®-RH96 Protein Analysis System (Sartorius, Göttingen, Germany). All experiments were performed at 30 °C with orbital agitation at 1,000 rpm.

To measure the affinity of PGT121 reference protein (LakePharma, San Carlos, California) and the PGT121 RibobNAb variants for BG505 DS-SOSIP.664 Env trimer (NIH), the antibodies were immobilized as ligands on the surface of Octet® Anti-hIgG Fc Capture (AHC) Biosensors (Sartorius, Göttingen, Germany), while soluble BG505 DS-SOSIP.664 (NIH) was used as analyte.

Prior to antibody immobilization, a baseline step using Octet® Kinetics Buffer (Sartorius, Göttingen, Germany) was performed for 60 s. Antibodies were immobilized at a concentration of 1 µg/mL for 900 s with a threshold of 1 nm, followed by a baseline step in Octet® Kinetics Buffer for 120 s. Binding affinities were determined using multi-cycle kinetics and decreasing SOSIP concentrations (26.71 nM to 0.42 nM, 2-fold dilutions in Octet® Kinetics Buffer). The association step was carried out for 600 s, followed by a 600 s dissociation step in Octet® Kinetics Buffer. In between cycles, the sensors were regenerated five times by incubating the sensors in glycine pH 1.5 buffer for 5 s, followed by a 5 s neutralization in Octet® Kinetics Buffer.

Data were analyzed using the Octet® Analysis Studio 13.0.1.35 software. To perform inter-step correction, the data were aligned with the dissociation step, and Savitzky-Golay filtering was applied to all curves. The association and dissociation curves were globally analyzed using a 1:1 Langmuir binding model.

### SOSIP-ELISA

A custom ELISA was performed to confirm binding of *in vitro*-expressed RibobNAbs to SOSIP. Streptavidin-coated 96-well plates (Thermo Fisher Scientific, Waltham, Massachusetts) were coated with 100 µL per well of biotinylated recombinant BG505 DS-SOSIP.664 Env trimer (ATUM, Newark, California; custom-made), at a concentration of 1 µg/mL in Coating Buffer (5 mM sodium carbonate, pH 9.6), and incubated overnight at 4 °C. Blocker™ Casein in PBS (Thermo Fisher Scientific, Waltham, Massachusetts) was diluted to 18% in PBS and used as blocking buffer. Plates were washed three times with PBS-T (phosphate-buffered saline containing 0.01% Tween-20) and blocked with 250 µL per well of blocking buffer for 1 h at 37 °C with gentle shaking. The samples, the PGT121 reference protein control (LakePharma, San Carlos, California) and an in-house produced negative control IgG were serially diluted in blocking buffer (ref. and neg. control: 800-0.0512 ng/mL; RNA-LNP transfection: 1:2, 1:6, 1:18,1:54, 1:162, 1:486, 1:1,458; RNA transfection: 1:5, 1:15, 1:45, 1:135, 1:405, 1:1,215, 1:3,645).

Following three washes with PBS-T, 100 µL of the diluted samples, reference and negative protein control were added to wells and incubated for 1 h at 37 °C with gentle shaking. After another three washing steps with PBS-T, 100 µL per well of HRP-conjugated goat anti-human IgG secondary antibody (Jackson ImmunoResearch Europe Ltd, Ely, England), diluted 1:5,000 in blocking buffer, was added and incubated for 45 min at 37 °C with gentle shaking. After washing three times with PBS-T, 100 µL per well of TMB substrate (Kementec Solutions A/S, Kuldyssen, Denmark) was added and incubated for 8 min at RT. The reaction was stopped with 100 µL per well of 25% sulphuric acid (Sigma-Aldrich, St. Louis, Missouri) and ΔOD_450-620 nm_ was quantified via BioTek Epoch Reader. RibobNAb concentrations in the diluted samples were determined using Gyros ELISA.

### Pseudovirus production

Pseudoviruses were generated following established protocols ^73^. HEK-293T-17 cells (50-80% confluent) were co-transfected with Env-expression plasmid and Env-deficient HIV-1 backbone vector (pSG3Δenv) using FuGENE 6 reagent according to the manufacturer’s instructions (FuGENE, Madison, Wisconsin). Following a 30-minute incubation period at RT (18 to 25 °C), transfection complexes were added to HEK-293T-17 cells. Cells were then incubated for 48 h with a single medium change performed 3 to 8 h post-transfection. Virus-containing supernatant was harvested from the culture flasks and passed through a 0.45 μm filter to remove cell debris. FBS was supplemented to achieve a final concentration of 20% in the virus-containing supernatant prior to storage at ≤-70 °C.

### TZM-bl pseudovirus neutralization assay

The TZM-bl neutralization assay was conducted following established protocols.^73^ Pseudoviruses, pre-titrated to yield luminescence signals of approximately 100,000 relative light units (RLU) or at least 10-fold above background levels, were combined with serial dilutions of RibobNAb-containing cell culture supernatants or NHP serum and co-incubated for 1 h at 37 °C. Subsequently, TZM-bl cells (10^4^ cells per well) were added to 96-well plates containing 250 µL of medium supplemented with 10-11 µg/mL DEAE-dextran. Control wells included cells with pseudovirus (without RibobNAb sample) and cells alone, serving as positive and negative infection controls, respectively. After 2 days of incubation at 37 °C with 5% CO_2_, luminescence detection was performed using a luminometer following the addition of Bright-Glo luciferase reagent (Promega, Madison, Wisconsin). Background RLU values from non-infected TZM-bl cells were subtracted, and IC_50_ and IC_80_ values were calculated as the antibody concentration resulting in a 50% or 80% RLU reduction, respectively, relative to untreated virus control wells.

### *In vivo* expression and pharmacokinetic studies in mice

The single-dose PK of PGT121 RibobNAb variants was evaluated in male and female NSG and NSG hFcRn Tg32 mice. NSG mice, aged 10 to 22 weeks, or NSG hFcRn mice, aged 6 to 10 weeks, were randomly distributed to test or control groups. Each group was further divided into two subgroups (“a” and “b”), with each subgroup consisting of three animals. For the single-dose PK assessment, mice in the test groups received 30 µg of the test items in 150 µL Dulbecco’s phosphate-buffered saline (DPBS), while the control group received 150 µL of DPBS alone. To generate serum samples, peripheral blood was collected at predetermined time points using Microvette 500 Z-Gel tubes (Sarstedt AG, Nümbrecht, Germany) with rotational sampling locations to minimize stress on individual animals. Blood sampling sites were rotated between the vena facialis, right retro-orbital sinus, and left retro-orbital sinus across different time points, with terminal exsanguination performed at study completion. The blood samples were centrifuged at 10,000 × g for 5 min at RT to separate the serum. Serum aliquots were stored at –65 to –85 °C until analysis by Gyros ELISA.

### Pharmacokinetic studies in NHPs

Five groups of rhesus monkeys (n = 3 non-naïve male animals per group) received either 1 mg/kg PGT121-LS reference protein administered IV, or PGT121-LS RNA-LNP at doses of 0.5 mg/kg or 1.6 mg/kg PGT121-LS delivered by single IV bolus or IM injections. IM groups were followed for a 120-day observation period and IV groups were monitored for a 180-day observation period. The dose volume was 3 mL for all animals, and the dose concentration was adjusted individually. The mean body weight was around 4.9 kg. Serial serum samples were collected at selected time points and stored at –70 °C until measurement. Serum concentrations were measured by a fit-for-purpose PK binding antibody multiplex assay (see below) to determine the amount of serum PGT121-LS IgG levels.

### PK-Binding antibody multiplex assay (BAMA)

PGT121-LS RibobNAb and reference protein levels in NHP serum were measured on a Bio-Plex instrument (Bio-Rad, Hercules, California) using a qualified assay designed to measure infused PGT121-LS RibobNAb/reference protein binding to an anti-idiotype antibody captured on fluorescent magnetic beads.^17^ This assay was derived from a standardized custom HIV-1 Luminex assay.^74–77^ The Bioplex software provides two readouts: a background-subtracted median fluorescent intensity (MFI), with background referring to a plate level control, and a concentration calculated using a standard curve with a 5PL curve fit. Concentrations are representative of three separate assays where each sample was run in duplicate. PGT121-LS was titrated in duplicate and replicate results were averaged to create a standard curve that was used to determine concentration of the diluted samples. The negative controls were CH58 (an irrelevant mAb) and blank beads. Samples with PGT121-LS concentrations below the lower limit of quantification (LLOQ) at a dilution of 1:100 were truncated at the LLOQ for plotting purposes. Samples with concentrations above the LLOQ at a 1:100 dilution were further tested at various dilution factors to obtain MFIs in the linear range of the standard curve, and the resulting concentrations from the 5PL standard curve were averaged. Samples that were positive (at or greater than the limit of detection) with an MFI that were 3-fold over the pre-infusion MFI with a concentration less than the LLOQ were called positive <LLOQ. The standard curve EC_50_ values and MFI values were tracked against historical data in Levey-Jennings and points with an MFI >100 must have had a %CV between replicates <20%. In addition, if the blank bead negative control exceeded 5,000 MFI, the sample was repeated. If the repeat value exceeded 5,000 MFI, the sample was excluded from analysis due to high background binding to magnetic beads.

### ADA assay

ADAs in serum from all 15 NHPs were detected and characterized using a tiered testing approach ^33,78^. Briefly, PGT121-LS was conjugated with either biotin or a sulfo-tag label. Biotinylated and sulfo-tagged antibodies (each at 50 ng/well) were then mixed an equal volume of serum (final serum dilution 50%), and ADAs were detected by bridging variably labeled forms of the drug. The level of biotinylated antibody bridged to sulfo-tagged PGT121-LS was measured using the Meso Scale Diagnostics™ platform (Meso Scale Discovery, Rockville, Maryland). A positivity cut point was established (Average signal + 3SD, i.e. 99.7^th^ percentile, from buffer only control) and utilized to determine whether a sample was ADA screening assay positive. Samples were tested in duplicate and percent coefficients of variation (%CVs) for all replicates were under 30%. Samples that were positive in screening were further titered by 3-fold dilution until no significant difference was observed between subsequent dilutions (Two-tailed Student’s t-test, α = p<0.05, Microsoft Excel 16.0300). The last significant dilution step determined the ADA titer (i.e. magnitude).

Each assay plate included technical blank samples (i.e. reagents without serum) and a positive control (i.e. anti-idiotype antibody 9E9-2 (mAb) Lot # 2017-12-1WS) at multiple test concentrations. Acceptance criteria were based on sample and control repeatability (%CV<30%) as well as positive controls retaining expected dose response and sensitivity (<100 ng/mL). ADA titers were plotted across time for all groups.

### Noncompartmental analysis (NCA)

Serum concentration-time data of reference protein or RNA-encoded PGT121-LS in mice and NHPs was evaluated by NCA using Phoenix WinNonlin, Version 8.3.5.340 (Certara, Princeton, NJ). AUC_last_ was calculated using the Linear Up Log Down method with extrapolated C_0_. AUC_inf_ was extrapolated using the predicted C_last_/λz. Profiles exhibiting an atypical PK behaviour (e.g. due to ADA development or rebound of concentrations at later time points), were excluded from calculations of mean values.

### Population PK model development

A two-compartment popPK model with linear elimination was developed for PGT121-LS in NHPs after single IV dosing (Figure 6A). Derived structural parameters including clearance (CL), intercompartmental clearance (Q), and central/peripheral distribution volumes (V_c_, V_p_) were subsequently fixed in the RNA-encoded PGT121-LS model, to estimate the RNA translation-related parameters (k_a_, F_max_ and FD_50_). PopPK modeling was conducted employing first-order conditional estimation extended least-squares (FOCE-ELS) algorithm using Phoenix NLME, Version 8.3.5.340 (Certara, Princeton, NJ). Non-linear translational efficiency was described using a saturable model:

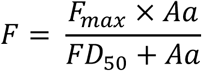

### Human PK prediction using allometric scaling

Human PK was predicted based on allometric scaling of PK parameters from NHPs. The allometric scaling exponents were assumed to be the same as published exponents for LS-engineered IgG (Table S4).^79^ RNA translation-related parameters including k_a_, F_max_ and FD_50_ were assumed to be identical between NHPs and humans.

Allometric scaling of human PK parameters was performed in Microsoft Excel according to the following formula:

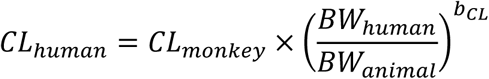

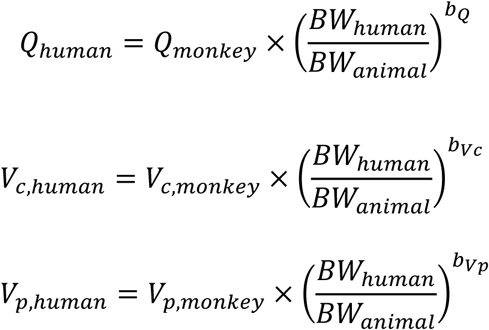

Abbreviations: b = Allometric exponent; BW = bodyweight; V_c_ = volume of distribution in the central compartment; V_p_ = volume of distribution in the peripheral compartment; CL = clearance from the central compartment; Q = intercompartmental clearance between central and peripheral compartment.

Using the predicted human PK parameters, PK profiles of reference protein or RNA-encoded PGT121-LS were simulated for single and repeated doses at 1 and 2 mg/kg every three months (Figure 6D and E).

## Data and materials availability

Requests for further information and resources should be directed to the lead contacts, Felix Tolksdorf and Valentin Le Douce. The PGT121 sequence has been reported before^15^ and can be obtained from GenBank (PGT121-VH: GenBank accession AEN14390.1 (amino acid), JN201894.1 (nucleotide); PGT121-VL: GenBank accession AEN14407.1 (amino acid), JN201911.1 (nucleotide)). Internally available proteins encoded by RNA-LNP samples or materials will be made available to academic, noncommercial researchers with a completed materials transfer agreement upon reasonable request. All data associated with this study can be found in the main text or the Supplementary Information. Source data are available upon reasonable request. This paper does not report original code. Any additional information required to reanalyze the data reported in this paper is available from the lead contact upon reasonable request.

## Acknowledgements

We thank Charles River Laboratories for NHP handling and non-clinical safety assessment, members of Georgia Tomaras’ laboratory for PK analysis, members of Margaret E. Ackerman’s laboratory for ADA analysis and Acuitas Therapeutics, Inc. for the LNP technology. We thank Mike McCune for reviewing the manuscript and Fiona Powell for editorial support. Illustrations were created with BioRender.com.

## Author contributions

F.T. and V.L.D. conceived the study; J.N. designed the RNA constructs; F.T. conceptualized the pharmacokinetic mouse studies; I.G. conducted all mouse experiments; R.J. conceived and conducted RNA and RNA-LNP transfections, Western blots and SOSIP-ELISA studies; L.F. conceived and A.M. conducted Gyros ELISA studies, J.P.B. conceived and C.M. conducted BLI binding studies; M.S.S. conceived the *in vitro* pVNT assay, A.C. designed the cynomolgus study; J.C. and M.K. performed pharmacokinetic analysis, data interpretation and PK modeling; J.A.W. and M.E.A. designed and oversaw ADA testing and reporting; N.L.Y. and G.D.T. designed and performed the PK assays; F.T., V.L.D., S.K., C.R.S., U.E. discussed the findings; U.Ş. initiated the project concept; A.K., F.T., V.L.D. and S.K. drafted the manuscript; A.K. drafted the final figures. All authors critically reviewed and approved the manuscript for submission.

## Competing interests

F.T., J.N., R.J., J.C., A.C., A.K., L.F., A.M., I.G., S.K., J.P.B., C.M., M.K., U.E., C.R.S. and U.Ş. are current employees of BioNTech SE. All authors own stock and/or stock options in BioNTech SE. U.Ş. is the chief executive officer of BioNTech SE. F.T, J.N. and V.L.D. are inventors on International Patent Application related to WO2023218431A1 and WO2025027585A1. S.K. is inventor on International Patent Application related to WO2025027585A1. G.D.T. is a scientific reviewer for the Gilead Scholars and received funding from her institution for other projects from BioNTech SE and UVax Bio LLC.

## Declaration of generative AI and AI-assisted technologies in the writing process

During the preparation of this work the authors used an internal AI tool based on GPT-5.1 and Claude-Sonnet 4 models to check grammar, spelling, punctuation and tone. After using this tool, the authors reviewed and edited the content as needed and take full responsibility for the content of the published article.

## List of supplementary materials

Figure S1 to S2

Table S1 to S4

## Funding

The work presented here was funded by BioNTech SE and the Gates Foundation (INV-029358). The conclusions and opinions expressed in this work are those of the author(s) alone and shall not be attributed to the Foundation. Under the grant conditions of the Foundation, a Creative Commons Attribution 4.0 License has already been assigned to the Author Accepted Manuscript version that might arise from this submission. Please note works submitted as a preprint have not undergone a peer review process.

