## Supplemental Data for "RNA-Encoded PGT121-LS Anti-HIV Antibody: Comprehensive Preclinical Characterization and Translational Pharmacokinetics"

### **Supplementary Information**

### **Supplementary Figures**

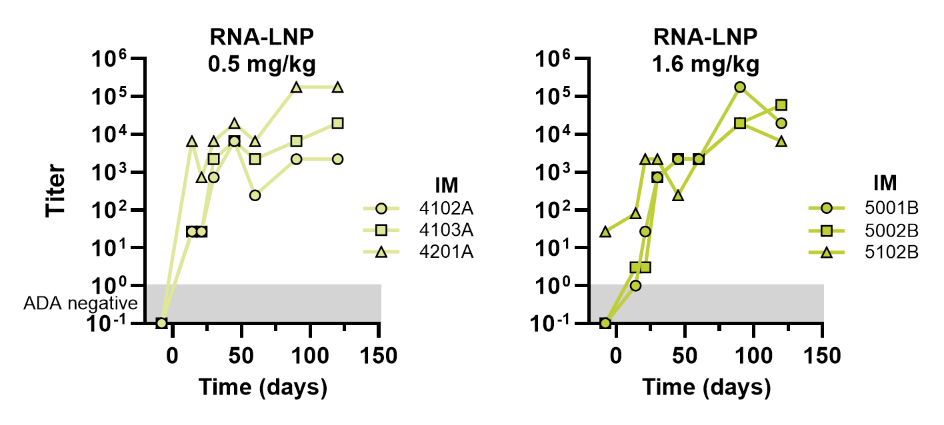

**Supplementary Figure 1:	ADA titers in NHP IM groups**

ADA positive samples were further titrated by 3-fold dilution until no significant difference was observed, with the last significant dilution step determining the ADA titer. Data for individual animals are shown.

ADA = anti-drug antibody; IM = intramuscular; LNP = lipid nanoparticle;RNA = ribonucleic acid.

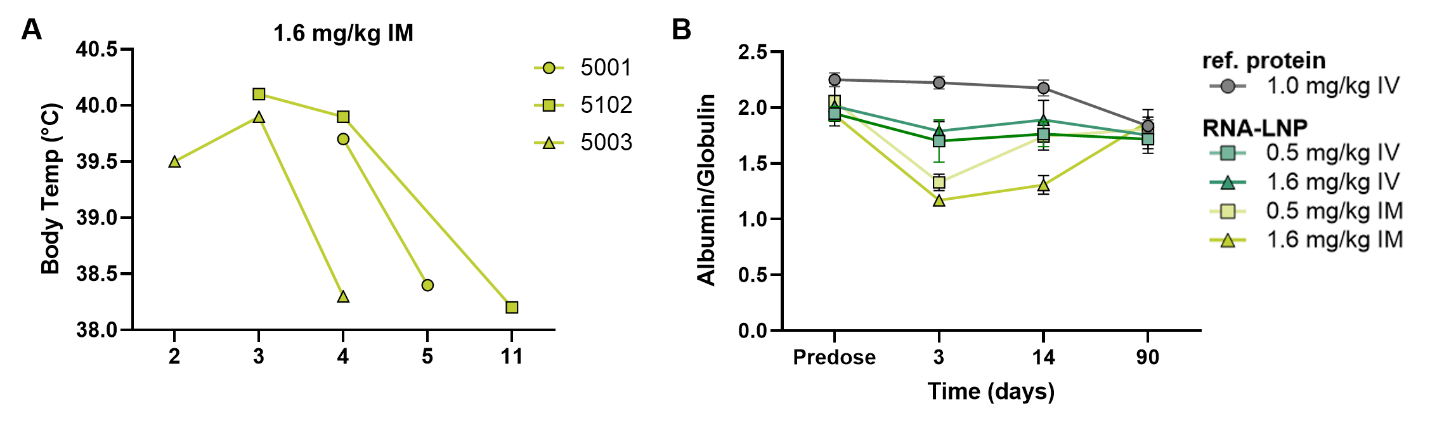

**Supplementary Figure 2:**	**Body temperatures and albumin/globulin ratios in NHPs**

A) Hyperthermia was noted in animals treated with 1.6 mg/kg PGT121-LS RNA-LNPs (animal nos. 5001, 5102 and 5003) from Day 2 to Day 4 with a return to normal body temperatures thereafter. B) Albumin/globulin ratios were determined predose and on Days 3, 14 and 90. IM injection of PGT121-LS RNA-LNPs resulted in a lower albumin/globulin ratio on Days 3 and 14.

IM = intramuscular; IV = intravenous; LNP = lipid nanoparticle; ref. = reference; RibobNAb = RNA-encoded, LNP-formulated anti-HIV-1 broadly neutralizing antibody; RNA = ribonucleic acid; Temp = temperature.

### **Supplementary Tables**

#### **Summary of hematology**

**Table S1**	**Summary of hematology I – Predose period**

| **GROUP** |  | **RBC** | **HGB** | **HCT** | **MCV** | **MCH** | **MCHC** | **RDW** | **PLT** |
| --- | --- | --- | --- | --- | --- | --- | --- | --- | --- |
| **NO.** |  | **(x10^12^/L)** | **(g/L)** | **(L/L)** | **(fL)** | **(pg)** | **(g/L)** | **(%)** | **(x10^9^/L)** |
| 1 | MEAN | 5.237 | 130.0 | 0.387 | 73.87 | 24.87 | 336.7 | 13.27 | 366.0 |
|  | SEM | 0.187 | 2.1 | 0.009 | 2.02 | 0.76 | 4.1 | 0.17 | 23.3 |
|  | N | 3 | 3 | 3 | 3 | 3 | 3 | 3 | 3 |
| 2 | MEAN | 5.677 | 135.3 | 0.410 | 72.43 | 23.83 | 329.3 | 13.10 | 428.3 |
|  | SEM | 0.126 | 3.5 | 0.010 | 0.44 | 0.38 | 3.3 | 0.36 | 64.3 |
|  | N | 3 | 3 | 3 | 3 | 3 | 3 | 3 | 3 |
| 3 | MEAN | 5.620 | 133.3 | 0.403 | 71.63 | 23.77 | 331.7 | 13.53 | 425.3 |
|  | SEM | 0.070 | 4.8 | 0.012 | 2.99 | 1.13 | 2.2 | 0.52 | 42.6 |
|  | N | 3 | 3 | 3 | 3 | 3 | 3 | 3 | 3 |
| 4 | MEAN | 5.380 | 133.7 | 0.397 | 74.00 | 24.87 | 336.7 | 13.67 | 316.0 |
|  | SEM | 0.225 | 4.1 | 0.007 | 2.38 | 0.23 | 7.9 | 0.66 | 12.0 |
|  | N | 3 | 3 | 3 | 3 | 3 | 3 | 3 | 3 |
| 5 | MEAN | 5.130 | 126.0 | 0.370 | 72.63 | 24.63 | 339.0 | 13.60 | 343.3 |
|  | SEM | 0.307 | 5.9 | 0.017 | 0.83 | 0.33 | 2.1 | 0.51 | 51.2 |
|  | N | 3 | 3 | 3 | 3 | 3 | 3 | 3 | 3 |

**Summary of hematology I – Day 3**

| **GROUP** |  | **RBC** | **HGB** | **HCT** | **MCV** | **MCH** | **MCHC** | **RDW** | **PLT** |
| --- | --- | --- | --- | --- | --- | --- | --- | --- | --- |
| **NO.** |  | **(x10^12^/L)** | **(g/L)** | **(L/L)** | **(fL)** | **(pg)** | **(g/L)** | **(%)** | **(x10^9^/L)** |
| 1 | MEAN | 4.820 | 114.0 | 0.370 | 76.43 | 23.77 | 311.0 | 13.93 | 366.0 |
|  | SEM | 0.319 | 4.2 | 0.015 | 2.55 | 0.82 | 1.0 | 0.12 | 33.6 |
|  | N | 3 | 3 | 3 | 3 | 3 | 3 | 3 | 3 |
| 2 | MEAN | 5.103 | 118.7 | 0.380 | 74.10 | 23.23 | 313.3 | 13.70 | 426.7 |
|  | SEM | 0.165 | 5.0 | 0.015 | 1.18 | 0.35 | 0.9 | 0.61 | 32.9 |
|  | N | 3 | 3 | 3 | 3 | 3 | 3 | 3 | 3 |
| 3 | MEAN | 5.107 | 118.7 | 0.380 | 74.53 | 23.23 | 312.0 | 14.00 | 226.7 |
|  | SEM | 0.038 | 5.2 | 0.015 | 3.26 | 1.16 | 3.2 | 0.40 | 105.0 |
|  | N | 3 | 3 | 3 | 3 | 3 | 3 | 3 | 3 |
| 4 | MEAN | 4.730 | 113.7 | 0.357 | 75.83 | 24.10 | 318.0 | 14.33 | 260.3 |
|  | SEM | 0.116 | 2.2 | 0.009 | 2.23 | 0.32 | 5.1 | 0.90 | 3.9 |
|  | N | 3 | 3 | 3 | 3 | 3 | 3 | 3 | 3 |
| 5 | MEAN | 4.913 | 121.0 | 0.373 | 75.67 | 24.67 | 326.0 | 13.63 | 279.7 |
|  | SEM | 0.344 | 5.8 | 0.020 | 1.15 | 0.49 | 2.0 | 0.43 | 40.5 |
|  | N | 3 | 3 | 3 | 3 | 3 | 3 | 3 | 3 |

**Summary of hematology I – Day 14**

| **GROUP** |  | **RBC** | **HGB** | **HCT** | **MCV** | **MCH** | **MCHC** | **RDW** | **PLT** |
| --- | --- | --- | --- | --- | --- | --- | --- | --- | --- |
| **NO.** |  | **(x10^12^/L)** | **(g/L)** | **(L/L)** | **(fL)** | **(pg)** | **(g/L)** | **(%)** | **(x10^9^/L)** |
| 1 | MEAN | 4.907 | 121.3 | 0.373 | 76.33 | 24.77 | 324.7 | 13.37 | 408.7 |
|  | SEM | 0.189 | 3.2 | 0.007 | 2.72 | 0.87 | 3.0 | 0.18 | 28.3 |
|  | N | 3 | 3 | 3 | 3 | 3 | 3 | 3 | 3 |
| 2 | MEAN | 5.363 | 126.7 | 0.400 | 74.23 | 23.63 | 318.0 | 13.50 | 471.3 |
|  | SEM | 0.111 | 4.8 | 0.015 | 1.47 | 0.48 | 1.2 | 0.35 | 40.8 |
|  | N | 3 | 3 | 3 | 3 | 3 | 3 | 3 | 3 |
| 3 | MEAN | 5.403 | 127.7 | 0.407 | 75.50 | 23.63 | 313.0 | 13.70 | 364.3 |
|  | SEM | 0.074 | 4.3 | 0.007 | 2.51 | 1.11 | 5.0 | 0.56 | 129.3 |
|  | N | 3 | 3 | 3 | 3 | 3 | 3 | 3 | 3 |
| 4 | MEAN | 5.067 | 123.7 | 0.390 | 77.27 | 24.40 | 316.7 | 14.53 | 387.7 |
|  | SEM | 0.216 | 3.3 | 0.006 | 2.60 | 0.42 | 5.5 | 0.71 | 16.9 |
|  | N | 3 | 3 | 3 | 3 | 3 | 3 | 3 | 3 |
| 5 | MEAN | 4.757 | 112.0 | 0.360 | 75.97 | 23.60 | 310.7 | 14.90 | 436.0 |
|  | SEM | 0.224 | 3.6 | 0.012 | 0.99 | 0.36 | 3.0 | 0.40 | 86.6 |
|  | N | 3 | 3 | 3 | 3 | 3 | 3 | 3 | 3 |

**Summary of hematology I – Day 30**

| **GROUP** |  | **RBC** | **HGB** | **HCT** | **MCV** | **MCH** | **MCHC** | **RDW** | **PLT** |
| --- | --- | --- | --- | --- | --- | --- | --- | --- | --- |
| **NO.** |  | **(x10^12^/L)** | **(g/L)** | **(L/L)** | **(fL)** | **(pg)** | **(g/L)** | **(%)** | **(x10^9^/L)** |
| 1 | MEAN | 5.013 | 123.7 | 0.377 | 75.87 | 24.77 | 326.7 | 12.97 | 369.7 |
|  | SEM | 0.258 | 3.5 | 0.009 | 3.63 | 0.97 | 2.8 | 0.22 | 40.4 |
|  | N | 3 | 3 | 3 | 3 | 3 | 3 | 3 | 3 |
| 2 | MEAN | 5.350 | 127.0 | 0.393 | 74.17 | 23.73 | 320.0 | 13.17 | 402.3 |
|  | SEM | 0.146 | 4.6 | 0.015 | 1.34 | 0.54 | 1.5 | 0.35 | 28.5 |
|  | N | 3 | 3 | 3 | 3 | 3 | 3 | 3 | 3 |
| 3 | MEAN | 5.440 | 128.7 | 0.407 | 74.90 | 23.63 | 315.7 | 13.17 | 431.0 |
|  | SEM | 0.050 | 6.0 | 0.019 | 3.20 | 1.05 | 2.0 | 0.41 | 66.5 |
|  | N | 3 | 3 | 3 | 3 | 3 | 3 | 3 | 3 |
| 4 | MEAN | 5.213 | 127.0 | 0.393 | 75.43 | 24.43 | 324.0 | 13.97 | 321.0 |
|  | SEM | 0.270 | 3.8 | 0.009 | 2.25 | 0.56 | 2.9 | 0.81 | 20.3 |
|  | N | 3 | 3 | 3 | 3 | 3 | 3 | 3 | 3 |
| 5 | MEAN | 5.163 | 125.3 | 0.390 | 75.57 | 24.30 | 322.3 | 13.43 | 295.3 |
|  | SEM | 0.272 | 5.8 | 0.015 | 0.96 | 0.17 | 1.9 | 0.20 | 53.1 |
|  | N | 3 | 3 | 3 | 3 | 3 | 3 | 3 | 3 |

**Summary of hematology I – Day 90**

| **GROUP** |  | **RBC** | **HGB** | **HCT** | **MCV** | **MCH** | **MCHC** | **RDW** | **PLT** |
| --- | --- | --- | --- | --- | --- | --- | --- | --- | --- |
| **NO.** |  | **(x10^12^/L)** | **(g/L)** | **(L/L)** | **(fL)** | **(pg)** | **(g/L)** | **(%)** | **(x10^9^/L)** |
| 1 | MEAN | 5.417 | 129.0 | 0.400 | 73.87 | 23.87 | 323.3 | 13.23 | 341.7 |
|  | SEM | 0.260 | 5.1 | 0.020 | 2.93 | 0.68 | 3.7 | 0.19 | 41.8 |
|  | N | 3 | 3 | 3 | 3 | 3 | 3 | 3 | 3 |
| 2 | MEAN | 5.753 | 131.7 | 0.410 | 71.63 | 22.90 | 319.7 | 13.13 | 377.7 |
|  | SEM | 0.283 | 6.3 | 0.020 | 1.07 | 0.64 | 4.1 | 0.20 | 44.8 |
|  | N | 3 | 3 | 3 | 3 | 3 | 3 | 3 | 3 |
| 3 | MEAN | 5.620 | 130.3 | 0.407 | 72.40 | 23.23 | 320.7 | 13.37 | 385.3 |
|  | SEM | 0.061 | 4.8 | 0.013 | 3.02 | 1.07 | 1.7 | 0.38 | 73.5 |
|  | N | 3 | 3 | 3 | 3 | 3 | 3 | 3 | 3 |
| 4 | MEAN | 5.720 | 135.0 | 0.423 | 73.90 | 23.67 | 321.0 | 14.37 | 298.7 |
|  | SEM | 0.229 | 2.9 | 0.003 | 2.64 | 0.47 | 5.5 | 0.57 | 16.8 |
|  | N | 3 | 3 | 3 | 3 | 3 | 3 | 3 | 3 |
| 5 | MEAN | 5.727 | 133.3 | 0.420 | 73.27 | 23.37 | 318.3 | 13.90 | 312.3 |
|  | SEM | 0.393 | 7.5 | 0.023 | 1.03 | 0.29 | 1.2 | 0.70 | 45.3 |
|  | N | 3 | 3 | 3 | 3 | 3 | 3 | 3 | 3 |

**Summary of hematology I – Day 129**

| **GROUP** |  | **RBC** | **HGB** | **HCT** | **MCV** | **MCH** | **MCHC** | **RDW** | **PLT** |
| --- | --- | --- | --- | --- | --- | --- | --- | --- | --- |
| **NO.** |  | **(x10^12^/L)** | **(g/L)** | **(L/L)** | **(fL)** | **(pg)** | **(g/L)** | **(%)** | **(x10^9^/L)** |
| 1 | MEAN | 5.220 | 127.3 | 0.387 | 74.17 | 24.43 | 330.0 | 13.90 | 405.0 |
|  | SEM | 0.226 | 4.1 | 0.009 | 2.72 | 0.52 | 4.9 | 0.10 | 53.2 |
|  | N | 3 | 3 | 3 | 3 | 3 | 3 | 3 | 3 |
| 2 | MEAN | 5.620 | 131.0 | 0.403 | 71.77 | 23.33 | 324.7 | 13.70 | 452.0 |
|  | SEM | 0.211 | 4.4 | 0.013 | 0.84 | 0.50 | 3.7 | 0.31 | 39.2 |
|  | N | 3 | 3 | 3 | 3 | 3 | 3 | 3 | 3 |
| 3 | MEAN | 5.347 | 126.3 | 0.390 | 73.17 | 23.73 | 324.3 | 14.07 | 498.3 |
|  | SEM | 0.149 | 2.9 | 0.006 | 3.09 | 1.18 | 2.8 | 0.47 | 53.7 |
|  | N | 3 | 3 | 3 | 3 | 3 | 3 | 3 | 3 |

**Summary of hematology I– Day 150**

| **GROUP** |  | **RBC** | **HGB** | **HCT** | **MCV** | **MCH** | **MCHC** | **RDW** | **PLT** |
| --- | --- | --- | --- | --- | --- | --- | --- | --- | --- |
| **NO.** |  | **(x10^12^/L)** | **(g/L)** | **(L/L)** | **(fL)** | **(pg)** | **(g/L)** | **(%)** | **(x10^9^/L)** |
| 1 | MEAN | 5.057 | 128.3 | 0.383 | 76.00 | 25.33 | 334.0 | 13.70 | 363.0 |
|  | SEM | 0.192 | 4.7 | 0.012 | 2.87 | 0.64 | 4.6 | 0.25 | 24.0 |
|  | N | 3 | 3 | 3 | 3 | 3 | 3 | 3 | 3 |
| 2 | MEAN | 5.483 | 132.3 | 0.403 | 73.97 | 24.10 | 325.3 | 13.57 | 406.3 |
|  | SEM | 0.150 | 3.5 | 0.009 | 1.36 | 0.58 | 2.2 | 0.39 | 35.5 |
|  | N | 3 | 3 | 3 | 3 | 3 | 3 | 3 | 3 |
| 3 | MEAN | 5.367 | 130.3 | 0.393 | 73.57 | 24.33 | 330.3 | 13.57 | 397.7 |
|  | SEM | 0.059 | 5.7 | 0.012 | 2.47 | 1.22 | 5.8 | 0.33 | 57.0 |
|  | N | 3 | 3 | 3 | 3 | 3 | 3 | 3 | 3 |

**Summary of hematology I – Day 180**

| **GROUP** |  | **RBC** | **HGB** | **HCT** | **MCV** | **MCH** | **MCHC** | **RDW** | **PLT** |
| --- | --- | --- | --- | --- | --- | --- | --- | --- | --- |
| **NO.** |  | **(x10^12^/L)** | **(g/L)** | **(L/L)** | **(fL)** | **(pg)** | **(g/L)** | **(%)** | **(x10^9^/L)** |
| 1 | MEAN | 5.240 | 131.7 | 0.403 | 76.93 | 25.20 | 328.0 | 13.17 | 358.3 |
|  | SEM | 0.146 | 1.9 | 0.009 | 2.88 | 0.67 | 3.5 | 0.12 | 32.5 |
|  | N | 3 | 3 | 3 | 3 | 3 | 3 | 3 | 3 |
| 2 | MEAN | 5.563 | 134.7 | 0.413 | 74.47 | 24.20 | 325.0 | 13.20 | 418.7 |
|  | SEM | 0.133 | 4.7 | 0.015 | 1.59 | 0.59 | 3.2 | 0.31 | 30.4 |
|  | N | 3 | 3 | 3 | 3 | 3 | 3 | 3 | 3 |
| 3 | MEAN | 5.387 | 131.7 | 0.403 | 74.37 | 24.53 | 329.3 | 13.23 | 398.3 |
|  | SEM | 0.101 | 4.4 | 0.007 | 2.73 | 1.26 | 5.2 | 0.42 | 37.1 |
|  | N | 3 | 3 | 3 | 3 | 3 | 3 | 3 | 3 |

HCT = hematocrit; HGB = hemoglobin; MCH = mean corpuscular hemoglobin; MCHC = mean corpuscular hemoglobin concentration; MCV = mean corpuscular volume; PLT = platelet count; RBC = red blood cell count; RDW = red cell distribution width.

**Summary of hematology II– Predose period**

| **GROUP** |  | **WBC** | **NEUT** | **LYMPH** | **MONO** | **EOS** | **BASO** | **LUC** |
| --- | --- | --- | --- | --- | --- | --- | --- | --- |
| **NO.** |  | **(x10^9^/L)** | **(x10^9^/L)** | **(x10^9^/L)** | **(x10^9^/L)** | **(x10^9^/L)** | **(x10^9^/L)** | **(x10^9^/L)** |
| 1 | MEAN | 9.867 | 1.707 | 7.613 | 0.323 | 0.073 | 0.050 | 0.097 |
|  | SEM | 1.984 | 0.130 | 1.737 | 0.067 | 0.022 | 0.012 | 0.029 |
|  | N | 3 | 3 | 3 | 3 | 3 | 3 | 3 |
| 2 | MEAN | 9.250 | 3.390 | 5.273 | 0.337 | 0.147 | 0.040 | 0.067 |
|  | SEM | 0.270 | 0.625 | 0.470 | 0.038 | 0.072 | 0.006 | 0.003 |
|  | N | 3 | 3 | 3 | 3 | 3 | 3 | 3 |
| 3 | MEAN | 8.020 | 2.677 | 4.867 | 0.263 | 0.093 | 0.037 | 0.080 |
|  | SEM | 0.919 | 0.642 | 1.383 | 0.044 | 0.030 | 0.009 | 0.021 |
|  | N | 3 | 3 | 3 | 3 | 3 | 3 | 3 |
| 4 | MEAN | 10.980 | 4.540 | 5.953 | 0.273 | 0.083 | 0.043 | 0.087 |
|  | SEM | 0.704 | 1.591 | 0.934 | 0.050 | 0.033 | 0.003 | 0.009 |
|  | N | 3 | 3 | 3 | 3 | 3 | 3 | 3 |
| 5 | MEAN | 7.937 | 3.750 | 3.677 | 0.320 | 0.093 | 0.027 | 0.067 |
|  | SEM | 1.210 | 1.160 | 0.714 | 0.042 | 0.056 | 0.007 | 0.007 |
|  | N | 3 | 3 | 3 | 3 | 3 | 3 | 3 |

**Summary of hematology II – Day 3**

| **GROUP** |  | **WBC** | **NEUT** | **LYMPH** | **MONO** | **EOS** | **BASO** | **LUC** |
| --- | --- | --- | --- | --- | --- | --- | --- | --- |
| **NO.** |  | **(x10^9^/L)** | **(x10^9^/L)** | **(x10^9^/L)** | **(x10^9^/L)** | **(x10^9^/L)** | **(x10^9^/L)** | **(x10^9^/L)** |
| 1 | MEAN | 10.193 | 3.947 | 5.673 | 0.287 | 0.090 | 0.087 | 0.107 |
|  | SEM | 1.901 | 1.307 | 0.962 | 0.067 | 0.055 | 0.024 | 0.023 |
|  | N | 3 | 3 | 3 | 3 | 3 | 3 | 3 |
| 2 | MEAN | 10.727 | 6.013 | 4.023 | 0.407 | 0.127 | 0.077 | 0.080 |
|  | SEM | 1.821 | 2.301 | 0.533 | 0.097 | 0.078 | 0.007 | 0.017 |
|  | N | 3 | 3 | 3 | 3 | 3 | 3 | 3 |
| 3 | MEAN | 9.877 | 5.077 | 4.180 | 0.347 | 0.097 | 0.067 | 0.107 |
|  | SEM | 2.613 | 1.934 | 0.950 | 0.143 | 0.027 | 0.012 | 0.042 |
|  | N | 3 | 3 | 3 | 3 | 3 | 3 | 3 |
| 4 | MEAN | 14.523 | 11.227 | 2.837 | 0.143 | 0.200 | 0.050 | 0.070 |
|  | SEM | 0.721 | 0.662 | 0.136 | 0.012 | 0.046 | 0.010 | 0.006 |
|  | N | 3 | 3 | 3 | 3 | 3 | 3 | 3 |
| 5 | MEAN | 12.317 | 9.800 | 2.093 | 0.130 | 0.220 | 0.030 | 0.053 |
|  | SEM | 1.850 | 2.101 | 0.180 | 0.021 | 0.101 | 0.006 | 0.007 |
|  | N | 3 | 3 | 3 | 3 | 3 | 3 | 3 |

**Summary of hematology II – Day 30**

| **GROUP** |  | **WBC** | **NEUT** | **LYMPH** | **MONO** | **EOS** | **BASO** | **LUC** |
| --- | --- | --- | --- | --- | --- | --- | --- | --- |
| **NO.** |  | **(x10^9^/L)** | **(x10^9^/L)** | **(x10^9^/L)** | **(x10^9^/L)** | **(x10^9^/L)** | **(x10^9^/L)** | **(x10^9^/L)** |
| 1 | MEAN | 13.233 | 3.977 | 8.507 | 0.487 | 0.083 | 0.063 | 0.117 |
|  | SEM | 1.506 | 1.190 | 1.859 | 0.107 | 0.032 | 0.018 | 0.033 |
|  | N | 3 | 3 | 3 | 3 | 3 | 3 | 3 |
| 2 | MEAN | 11.833 | 4.783 | 6.123 | 0.430 | 0.357 | 0.057 | 0.080 |
|  | SEM | 2.229 | 0.991 | 1.038 | 0.070 | 0.106 | 0.012 | 0.017 |
|  | N | 3 | 3 | 3 | 3 | 3 | 3 | 3 |
| 3 | MEAN | 10.003 | 2.627 | 6.607 | 0.420 | 0.183 | 0.047 | 0.120 |
|  | SEM | 0.737 | 0.593 | 1.039 | 0.084 | 0.047 | 0.007 | 0.017 |
|  | N | 3 | 3 | 3 | 3 | 3 | 3 | 3 |
| 4 | MEAN | 10.773 | 2.407 | 7.737 | 0.380 | 0.133 | 0.050 | 0.070 |
|  | SEM | 1.107 | 0.175 | 1.026 | 0.051 | 0.026 | 0.006 | 0.010 |
|  | N | 3 | 3 | 3 | 3 | 3 | 3 | 3 |
| 5 | MEAN | 9.280 | 3.513 | 5.143 | 0.440 | 0.077 | 0.040 | 0.067 |
|  | SEM | 1.422 | 1.089 | 0.425 | 0.079 | 0.022 | 0.006 | 0.003 |
|  | N | 3 | 3 | 3 | 3 | 3 | 3 | 3 |

**Summary of hematology II – Day 90**

| **GROUP** |  | **WBC** | **NEUT** | **LYMPH** | **MONO** | **EOS** | **BASO** | **LUC** |
| --- | --- | --- | --- | --- | --- | --- | --- | --- |
| **NO.** |  | **(x10^9^/L)** | **(x10^9^/L)** | **(x10^9^/L)** | **(x10^9^/L)** | **(x10^9^/L)** | **(x10^9^/L)** | **(x10^9^/L)** |
| 1 | MEAN | 13.247 | 3.410 | 9.177 | 0.377 | 0.103 | 0.063 | 0.110 |
|  | SEM | 2.335 | 0.664 | 2.353 | 0.050 | 0.037 | 0.020 | 0.036 |
|  | N | 3 | 3 | 3 | 3 | 3 | 3 | 3 |
| 2 | MEAN | 9.673 | 1.863 | 6.957 | 0.417 | 0.277 | 0.050 | 0.113 |
|  | SEM | 0.839 | 0.535 | 0.445 | 0.118 | 0.097 | 0.006 | 0.034 |
|  | N | 3 | 3 | 3 | 3 | 3 | 3 | 3 |
| 3 | MEAN | 10.397 | 3.833 | 5.987 | 0.313 | 0.123 | 0.063 | 0.073 |
|  | SEM | 2.403 | 1.777 | 0.913 | 0.067 | 0.047 | 0.020 | 0.030 |
|  | N | 3 | 3 | 3 | 3 | 3 | 3 | 3 |
| 4 | MEAN | 11.903 | 3.033 | 8.220 | 0.347 | 0.133 | 0.063 | 0.103 |
|  | SEM | 1.103 | 0.273 | 0.798 | 0.085 | 0.032 | 0.007 | 0.023 |
|  | N | 3 | 3 | 3 | 3 | 3 | 3 | 3 |
| 5 | MEAN | 11.527 | 4.193 | 6.447 | 0.507 | 0.220 | 0.073 | 0.090 |
|  | SEM | 2.670 | 2.658 | 0.194 | 0.121 | 0.074 | 0.012 | 0.006 |
|  | N | 3 | 3 | 3 | 3 | 3 | 3 | 3 |

**Summary of hematology II – Day 129**

| **GROUP** |  | **WBC** | **NEUT** | **LYMPH** | **MONO** | **EOS** | **BASO** | **LUC** |
| --- | --- | --- | --- | --- | --- | --- | --- | --- |
| **NO.** |  | **(x10^9^/L)** | **(x10^9^/L)** | **(x10^9^/L)** | **(x10^9^/L)** | **(x10^9^/L)** | **(x10^9^/L)** | **(x10^9^/L)** |
| 1 | MEAN | 10.340 | 2.483 | 7.227 | 0.360 | 0.093 | 0.063 | 0.110 |
|  | SEM | 2.420 | 0.523 | 1.695 | 0.098 | 0.050 | 0.020 | 0.040 |
|  | N | 3 | 3 | 3 | 3 | 3 | 3 | 3 |
| 2 | MEAN | 10.273 | 2.640 | 6.827 | 0.390 | 0.280 | 0.060 | 0.083 |
|  | SEM | 1.361 | 0.643 | 0.700 | 0.055 | 0.121 | 0.015 | 0.012 |
|  | N | 3 | 3 | 3 | 3 | 3 | 3 | 3 |
| 3 | MEAN | 11.580 | 4.323 | 6.453 | 0.407 | 0.210 | 0.063 | 0.113 |
|  | SEM | 3.101 | 1.744 | 1.440 | 0.097 | 0.075 | 0.013 | 0.035 |
|  | N | 3 | 3 | 3 | 3 | 3 | 3 | 3 |

**Summary of hematology II – Day 150**

| **GROUP** |  | **WBC** | **NEUT** | **LYMPH** | **MONO** | **EOS** | **BASO** | **LUC** |
| --- | --- | --- | --- | --- | --- | --- | --- | --- |
| **NO.** |  | **(x10^9^/L)** | **(x10^9^/L)** | **(x10^9^/L)** | **(x10^9^/L)** | **(x10^9^/L)** | **(x10^9^/L)** | **(x10^9^/L)** |
| 1 | MEAN | 11.357 | 4.977 | 5.730 | 0.343 | 0.083 | 0.110 | 0.110 |
|  | SEM | 3.494 | 2.863 | 1.030 | 0.140 | 0.063 | 0.029 | 0.029 |
|  | N | 3 | 3 | 3 | 3 | 3 | 3 | 3 |
| 2 | MEAN | 9.493 | 2.433 | 6.327 | 0.343 | 0.167 | 0.127 | 0.090 |
|  | SEM | 1.902 | 0.837 | 1.009 | 0.102 | 0.038 | 0.032 | 0.015 |
|  | N | 3 | 3 | 3 | 3 | 3 | 3 | 3 |
| 3 | MEAN | 9.470 | 2.983 | 5.840 | 0.237 | 0.187 | 0.130 | 0.093 |
|  | SEM | 2.463 | 0.888 | 1.457 | 0.057 | 0.073 | 0.046 | 0.023 |
|  | N | 3 | 3 | 3 | 3 | 3 | 3 | 3 |

**Summary of hematology II – Day 180**

| **GROUP** |  | **WBC** | **NEUT** | **LYMPH** | **MONO** | **EOS** | **BASO** | **LUC** |
| --- | --- | --- | --- | --- | --- | --- | --- | --- |
| **NO.** |  | **(x10^9^/L)** | **(x10^9^/L)** | **(x10^9^/L)** | **(x10^9^/L)** | **(x10^9^/L)** | **(x10^9^/L)** | **(x10^9^/L)** |
| 1 | MEAN | 10.943 | 2.183 | 8.053 | 0.410 | 0.127 | 0.053 | 0.117 |
|  | SEM | 2.190 | 0.079 | 1.878 | 0.136 | 0.055 | 0.020 | 0.041 |
|  | N | 3 | 3 | 3 | 3 | 3 | 3 | 3 |
| 2 | MEAN | 8.820 | 2.050 | 6.157 | 0.353 | 0.180 | 0.030 | 0.053 |
|  | SEM | 1.730 | 0.680 | 1.040 | 0.083 | 0.012 | 0.006 | 0.018 |
|  | N | 3 | 3 | 3 | 3 | 3 | 3 | 3 |
| 3 | MEAN | 9.557 | 2.490 | 6.453 | 0.337 | 0.163 | 0.040 | 0.077 |
|  | SEM | 1.066 | 0.674 | 1.547 | 0.072 | 0.058 | 0.010 | 0.023 |
|  | N | 3 | 3 | 3 | 3 | 3 | 3 | 3 |

BASO = basophils;EOS = eosinophils; LUC = large unstained cells; LYMPH = lymphocytes; MONO = monocytes; NEUT = neutrophils; WBC = white blood cell count.

**Summary of hematology III – Predose period**

| **GROUP** |  | **NEUT** | **LYMPH** | **MONO** | **EOS** | **BASO** | **LUC** | **RETIC** | **RETIC** |
| --- | --- | --- | --- | --- | --- | --- | --- | --- | --- |
| **NO.** |  | **(%)** | **(%)** | **(%)** | **(%)** | **(%)** | **(%)** | **(x10^9^/L)** | **(%)** |
| 1 | MEAN | 18.43 | 76.07 | 3.37 | 0.70 | 0.50 | 1.00 | 56.57 | 1.07 |
|  | SEM | 2.85 | 2.84 | 0.28 | 0.17 | 0.06 | 0.10 | 3.85 | 0.07 |
|  | N | 3 | 3 | 3 | 3 | 3 | 3 | 3 | 3 |
| 2 | MEAN | 36.37 | 57.37 | 3.57 | 1.57 | 0.40 | 0.77 | 64.83 | 1.17 |
|  | SEM | 6.01 | 6.60 | 0.33 | 0.78 | 0.06 | 0.03 | 4.93 | 0.09 |
|  | N | 3 | 3 | 3 | 3 | 3 | 3 | 3 | 3 |
| 3 | MEAN | 35.83 | 58.37 | 3.23 | 1.17 | 0.47 | 0.97 | 55.70 | 1.00 |
|  | SEM | 11.64 | 11.45 | 0.28 | 0.27 | 0.09 | 0.15 | 8.25 | 0.17 |
|  | N | 3 | 3 | 3 | 3 | 3 | 3 | 3 | 3 |
| 4 | MEAN | 39.97 | 55.60 | 2.47 | 0.80 | 0.43 | 0.77 | 51.13 | 1.00 |
|  | SEM | 11.46 | 11.17 | 0.34 | 0.31 | 0.07 | 0.13 | 7.07 | 0.17 |
|  | N | 3 | 3 | 3 | 3 | 3 | 3 | 3 | 3 |
| 5 | MEAN | 45.43 | 48.27 | 4.03 | 1.10 | 0.33 | 0.83 | 53.07 | 1.03 |
|  | SEM | 11.60 | 11.23 | 0.12 | 0.55 | 0.07 | 0.09 | 5.56 | 0.03 |
|  | N | 3 | 3 | 3 | 3 | 3 | 3 | 3 | 3 |

**Summary of hematology III – Day 3**

| **GROUP** |  | **NEUT** | **LYMPH** | **MONO** | **EOS** | **BASO** | **LUC** | **RETIC** | **RETIC** |
| --- | --- | --- | --- | --- | --- | --- | --- | --- | --- |
| **NO.** |  | **(%)** | **(%)** | **(%)** | **(%)** | **(%)** | **(%)** | **(x10^9^/L)** | **(%)** |
| 1 | MEAN | 37.80 | 56.57 | 2.80 | 0.93 | 0.83 | 1.00 | 81.50 | 1.73 |
|  | SEM | 7.61 | 7.19 | 0.31 | 0.58 | 0.15 | 0.06 | 8.18 | 0.27 |
|  | N | 3 | 3 | 3 | 3 | 3 | 3 | 3 | 3 |
| 2 | MEAN | 51.57 | 41.87 | 3.87 | 1.23 | 0.77 | 0.77 | 109.93 | 2.13 |
|  | SEM | 13.22 | 12.86 | 0.76 | 0.67 | 0.07 | 0.12 | 20.23 | 0.39 |
|  | N | 3 | 3 | 3 | 3 | 3 | 3 | 3 | 3 |
| 3 | MEAN | 49.07 | 44.90 | 3.30 | 0.97 | 0.70 | 1.07 | 94.60 | 1.87 |
|  | SEM | 10.62 | 11.53 | 0.80 | 0.09 | 0.06 | 0.12 | 14.17 | 0.28 |
|  | N | 3 | 3 | 3 | 3 | 3 | 3 | 3 | 3 |
| 4 | MEAN | 77.23 | 19.57 | 1.00 | 1.37 | 0.33 | 0.47 | 60.27 | 1.27 |
|  | SEM | 1.15 | 0.76 | 0.10 | 0.38 | 0.03 | 0.09 | 0.29 | 0.03 |
|  | N | 3 | 3 | 3 | 3 | 3 | 3 | 3 | 3 |
| 5 | MEAN | 78.13 | 18.07 | 1.07 | 2.03 | 0.23 | 0.43 | 46.07 | 0.90 |
|  | SEM | 4.81 | 3.77 | 0.09 | 0.97 | 0.09 | 0.09 | 10.97 | 0.15 |
|  | N | 3 | 3 | 3 | 3 | 3 | 3 | 3 | 3 |

**Summary of hematology III – Day 14**

| **GROUP** |  | **NEUT** | **LYMPH** | **MONO** | **EOS** | **BASO** | **LUC** | **RETIC** | **RETIC** |
| --- | --- | --- | --- | --- | --- | --- | --- | --- | --- |
| **NO.** |  | **(%)** | **(%)** | **(%)** | **(%)** | **(%)** | **(%)** | **(x10^9^/L)** | **(%)** |
| 1 | MEAN | 23.00 | 71.23 | 3.23 | 1.00 | 0.50 | 1.03 | 100.83 | 2.07 |
|  | SEM | 2.80 | 2.70 | 0.38 | 0.21 | 0.06 | 0.15 | 14.75 | 0.32 |
|  | N | 3 | 3 | 3 | 3 | 3 | 3 | 3 | 3 |
| 2 | MEAN | 24.37 | 67.77 | 3.73 | 2.77 | 0.50 | 0.87 | 112.80 | 2.13 |
|  | SEM | 8.02 | 7.00 | 0.24 | 1.07 | 0.10 | 0.07 | 5.26 | 0.09 |
|  | N | 3 | 3 | 3 | 3 | 3 | 3 | 3 | 3 |
| 3 | MEAN | 36.13 | 57.33 | 3.40 | 1.43 | 0.47 | 1.23 | 99.50 | 1.87 |
|  | SEM | 8.33 | 8.37 | 0.40 | 0.55 | 0.03 | 0.15 | 14.02 | 0.28 |
|  | N | 3 | 3 | 3 | 3 | 3 | 3 | 3 | 3 |
| 4 | MEAN | 42.60 | 51.93 | 3.07 | 1.30 | 0.47 | 0.70 | 105.10 | 2.10 |
|  | SEM | 1.59 | 2.04 | 0.78 | 0.25 | 0.03 | 0.06 | 9.15 | 0.17 |
|  | N | 3 | 3 | 3 | 3 | 3 | 3 | 3 | 3 |
| 5 | MEAN | 27.77 | 63.00 | 3.37 | 4.00 | 1.00 | 0.87 | 151.77 | 3.20 |
|  | SEM | 6.39 | 5.28 | 0.53 | 1.30 | 0.06 | 0.03 | 12.94 | 0.21 |
|  | N | 3 | 3 | 3 | 3 | 3 | 3 | 3 | 3 |

**Summary of hematology III – Day 30**

| **GROUP** |  | **NEUT** | **LYMPH** | **MONO** | **EOS** | **BASO** | **LUC** | **RETIC** | **RETIC** |
| --- | --- | --- | --- | --- | --- | --- | --- | --- | --- |
| **NO.** |  | **(%)** | **(%)** | **(%)** | **(%)** | **(%)** | **(%)** | **(x10^9^/L)** | **(%)** |
| 1 | MEAN | 30.97 | 63.50 | 3.60 | 0.63 | 0.40 | 0.83 | 71.27 | 1.43 |
|  | SEM | 10.98 | 10.55 | 0.38 | 0.23 | 0.10 | 0.15 | 5.23 | 0.18 |
|  | N | 3 | 3 | 3 | 3 | 3 | 3 | 3 | 3 |
| 2 | MEAN | 40.03 | 52.27 | 3.70 | 2.87 | 0.47 | 0.67 | 85.97 | 1.63 |
|  | SEM | 1.07 | 1.32 | 0.15 | 0.43 | 0.09 | 0.03 | 5.66 | 0.12 |
|  | N | 3 | 3 | 3 | 3 | 3 | 3 | 3 | 3 |
| 3 | MEAN | 26.83 | 65.53 | 4.17 | 1.83 | 0.47 | 1.17 | 69.60 | 1.27 |
|  | SEM | 7.20 | 8.03 | 0.68 | 0.48 | 0.03 | 0.09 | 12.12 | 0.23 |
|  | N | 3 | 3 | 3 | 3 | 3 | 3 | 3 | 3 |
| 4 | MEAN | 22.53 | 71.30 | 3.67 | 1.33 | 0.43 | 0.67 | 64.53 | 1.27 |
|  | SEM | 1.58 | 2.40 | 0.82 | 0.39 | 0.03 | 0.03 | 3.34 | 0.12 |
|  | N | 3 | 3 | 3 | 3 | 3 | 3 | 3 | 3 |
| 5 | MEAN | 35.80 | 57.43 | 4.77 | 0.80 | 0.43 | 0.73 | 52.80 | 1.00 |
|  | SEM | 8.01 | 7.77 | 0.24 | 0.15 | 0.03 | 0.13 | 13.68 | 0.21 |
|  | N | 3 | 3 | 3 | 3 | 3 | 3 | 3 | 3 |

**Summary of hematology III – Day 90**

| **GROUP** |  | **NEUT** | **LYMPH** | **MONO** | **EOS** | **BASO** | **LUC** | **RETIC** | **RETIC** |
| --- | --- | --- | --- | --- | --- | --- | --- | --- | --- |
| **NO.** |  | **(%)** | **(%)** | **(%)** | **(%)** | **(%)** | **(%)** | **(x10^9^/L)** | **(%)** |
| 1 | MEAN | 27.50 | 67.53 | 2.93 | 0.73 | 0.43 | 0.80 | 45.80 | 0.83 |
|  | SEM | 8.11 | 8.23 | 0.48 | 0.23 | 0.07 | 0.12 | 6.11 | 0.13 |
|  | N | 3 | 3 | 3 | 3 | 3 | 3 | 3 | 3 |
| 2 | MEAN | 18.63 | 72.23 | 4.37 | 2.97 | 0.53 | 1.23 | 47.23 | 0.83 |
|  | SEM | 3.64 | 2.62 | 1.48 | 1.13 | 0.07 | 0.43 | 6.73 | 0.15 |
|  | N | 3 | 3 | 3 | 3 | 3 | 3 | 3 | 3 |
| 3 | MEAN | 33.93 | 60.30 | 3.10 | 1.40 | 0.60 | 0.70 | 44.23 | 0.80 |
|  | SEM | 8.49 | 8.81 | 0.53 | 0.75 | 0.06 | 0.12 | 12.40 | 0.21 |
|  | N | 3 | 3 | 3 | 3 | 3 | 3 | 3 | 3 |
| 4 | MEAN | 25.53 | 69.10 | 2.90 | 1.07 | 0.57 | 0.83 | 34.03 | 0.60 |
|  | SEM | 1.08 | 1.89 | 0.70 | 0.24 | 0.09 | 0.09 | 2.35 | 0.06 |
|  | N | 3 | 3 | 3 | 3 | 3 | 3 | 3 | 3 |
| 5 | MEAN | 30.07 | 62.07 | 4.43 | 2.00 | 0.67 | 0.83 | 40.60 | 0.73 |
|  | SEM | 13.60 | 13.25 | 0.75 | 0.69 | 0.12 | 0.12 | 4.72 | 0.03 |
|  | N | 3 | 3 | 3 | 3 | 3 | 3 | 3 | 3 |

**Summary of hematology III – Day 129**

| **GROUP** |  | **NEUT** | **LYMPH** | **MONO** | **EOS** | **BASO** | **LUC** | **RETIC** | **RETIC** |
| --- | --- | --- | --- | --- | --- | --- | --- | --- | --- |
| **NO.** |  | **(%)** | **(%)** | **(%)** | **(%)** | **(%)** | **(%)** | **(x10^9^/L)** | **(%)** |
| 1 | MEAN | 24.33 | 69.90 | 3.43 | 0.77 | 0.57 | 1.03 | 62.00 | 1.17 |
|  | SEM | 0.62 | 0.12 | 0.23 | 0.29 | 0.09 | 0.15 | 7.20 | 0.19 |
|  | N | 3 | 3 | 3 | 3 | 3 | 3 | 3 | 3 |
| 2 | MEAN | 25.13 | 67.10 | 3.77 | 2.70 | 0.60 | 0.77 | 72.83 | 1.30 |
|  | SEM | 3.00 | 3.84 | 0.09 | 1.01 | 0.10 | 0.03 | 4.27 | 0.12 |
|  | N | 3 | 3 | 3 | 3 | 3 | 3 | 3 | 3 |
| 3 | MEAN | 35.37 | 57.47 | 3.57 | 2.07 | 0.60 | 1.00 | 80.47 | 1.53 |
|  | SEM | 7.19 | 7.84 | 0.43 | 0.97 | 0.06 | 0.23 | 15.57 | 0.33 |
|  | N | 3 | 3 | 3 | 3 | 3 | 3 | 3 | 3 |

**Summary of hematology III – Day 150**

| **GROUP** |  | **NEUT** | **LYMPH** | **MONO** | **EOS** | **BASO** | **LUC** | **RETIC** | **RETIC** |
| --- | --- | --- | --- | --- | --- | --- | --- | --- | --- |
| **NO.** |  | **(%)** | **(%)** | **(%)** | **(%)** | **(%)** | **(%)** | **(x10^9^/L)** | **(%)** |
| 1 | MEAN | 38.47 | 55.80 | 2.83 | 0.90 | 1.03 | 1.00 | 64.07 | 1.23 |
|  | SEM | 11.31 | 10.91 | 0.30 | 0.66 | 0.09 | 0.06 | 2.95 | 0.03 |
|  | N | 3 | 3 | 3 | 3 | 3 | 3 | 3 | 3 |
| 2 | MEAN | 24.37 | 67.93 | 3.47 | 1.97 | 1.30 | 1.00 | 88.60 | 1.63 |
|  | SEM | 3.87 | 3.46 | 0.62 | 0.53 | 0.06 | 0.12 | 12.82 | 0.30 |
|  | N | 3 | 3 | 3 | 3 | 3 | 3 | 3 | 3 |
| 3 | MEAN | 31.00 | 61.83 | 2.57 | 2.27 | 1.30 | 1.00 | 57.53 | 1.07 |
|  | SEM | 1.27 | 2.34 | 0.35 | 1.14 | 0.10 | 0.06 | 5.34 | 0.09 |
|  | N | 3 | 3 | 3 | 3 | 3 | 3 | 3 | 3 |

**Summary of hematology III – Day 180**

| **GROUP** |  | **NEUT** | **LYMPH** | **MONO** | **EOS** | **BASO** | **LUC** | **RETIC** | **RETIC** |
| --- | --- | --- | --- | --- | --- | --- | --- | --- | --- |
| **NO.** |  | **(%)** | **(%)** | **(%)** | **(%)** | **(%)** | **(%)** | **(x10^9^/L)** | **(%)** |
| 1 | MEAN | 21.63 | 72.30 | 3.60 | 1.03 | 0.47 | 0.97 | 55.43 | 1.07 |
|  | SEM | 4.20 | 3.35 | 0.55 | 0.37 | 0.09 | 0.18 | 5.83 | 0.09 |
|  | N | 3 | 3 | 3 | 3 | 3 | 3 | 3 | 3 |
| 2 | MEAN | 22.43 | 70.47 | 3.97 | 2.23 | 0.30 | 0.60 | 61.60 | 1.10 |
|  | SEM | 3.49 | 2.92 | 0.50 | 0.55 | 0.00 | 0.21 | 1.58 | 0.06 |
|  | N | 3 | 3 | 3 | 3 | 3 | 3 | 3 | 3 |
| 3 | MEAN | 27.77 | 65.87 | 3.43 | 1.73 | 0.40 | 0.77 | 49.97 | 0.93 |
|  | SEM | 9.70 | 10.04 | 0.35 | 0.71 | 0.06 | 0.18 | 8.28 | 0.18 |
|  | N | 3 | 3 | 3 | 3 | 3 | 3 | 3 | 3 |

BASO = basophils; EOS = eosinophils; LUC = large unstained cells; LYMPH = lymphocytes; MONO = monocytes; NEUT = neutrophils; RETIC = reticulocytes.

#### **Summary of clinical chemistry**

**Table S2:**	**Summary of clinical chemistry I – Predose**

| **GROUP** |  | **AST** | **ALT** | **ALP** | **TBIL** | **CHOL** | **TRIG** | **GLUC** | **UREA** | **CREAT** |
| --- | --- | --- | --- | --- | --- | --- | --- | --- | --- | --- |
| **NO.** |  | **(U/L)** | **(U/L)** | **(U/L)** | **(μmol/L)** | **(mmol/L)** | **(mmol/L)** | **(mmol/L)** | **(mmol/L)** | **(μmol/L)** |
| 1 | MEAN | 28.0 | 42.3 | 514.7 | 3.15 | 3.360 | 0.797 | 5.57 | 5.70 | 58.0 |
|  | SEM | 2.3 | 7.4 | 39.0 | 0.05 | 0.236 | 0.114 | 0.03 | 0.36 | 2.3 |
|  | N | 3 | 3 | 3 | 2 | 3 | 3 | 3 | 3 | 3 |
| 2 | MEAN | 34.0 | 37.3 | 560.7 | 2.90 | 2.847 | 0.560 | 4.47 | 6.33 | 76.0 |
|  | SEM | 2.0 | 2.2 | 91.7 | 0.40 | 0.311 | 0.081 | 0.30 | 0.42 | 10.7 |
|  | N | 3 | 3 | 3 | 3 | 3 | 3 | 3 | 3 | 3 |
| 3 | MEAN | 33.7 | 34.3 | 544.0 | 4.20 | 2.840 | 0.667 | 4.73 | 5.43 | 67.7 |
|  | SEM | 2.0 | 2.7 | 141.7 | 1.30 | 0.283 | 0.151 | 0.38 | 0.37 | 3.7 |
|  | N | 3 | 3 | 3 | 2 | 3 | 3 | 3 | 3 | 3 |
| 4 | MEAN | 28.0 | 31.0 | 505.7 | 3.00 | 3.003 | 0.483 | 4.77 | 5.60 | 61.0 |
|  | SEM | 3.2 | 1.7 | 22.8 | 0.10 | 0.217 | 0.038 | 0.68 | 0.35 | 1.7 |
|  | N | 3 | 3 | 3 | 2 | 3 | 3 | 3 | 3 | 3 |
| 5 | MEAN | 31.3 | 37.0 | 504.3 | 3.13 | 3.473 | 0.460 | 3.70 | 5.47 | 49.3 |
|  | SEM | 3.5 | 7.8 | 152.3 | 0.53 | 0.237 | 0.074 | 0.25 | 0.28 | 2.9 |
|  | N | 3 | 3 | 3 | 3 | 3 | 3 | 3 | 3 | 3 |

**Summary of clinical chemistry I – Day 3**

| **GROUP** |  | **AST** | **ALT** | **ALP** | **TBIL** | **CHOL** | **TRIG** | **GLUC** | **UREA** | **CREAT** |
| --- | --- | --- | --- | --- | --- | --- | --- | --- | --- | --- |
| **NO.** |  | **(U/L)** | **(U/L)** | **(U/L)** | **(μmol/L)** | **(mmol/L)** | **(mmol/L)** | **(mmol/L)** | **(mmol/L)** | **(μmol/L)** |
| 1 | MEAN | 28.3 | 46.7 | 557.7 | 3.80 | 3.177 | 0.697 | 4.67 | 5.40 | 58.7 |
|  | SEM | 1.2 | 7.8 | 44.5 | 0.61 | 0.274 | 0.225 | 0.23 | 0.10 | 2.7 |
|  | N | 3 | 3 | 3 | 3 | 3 | 3 | 3 | 3 | 3 |
| 2 | MEAN | 42.7 | 54.7 | 577.3 | 3.60 | 2.563 | 0.660 | 4.10 | 7.07 | 74.7 |
|  | SEM | 10.2 | 8.4 | 77.7 | 0.20 | 0.280 | 0.106 | 0.38 | 0.52 | 6.9 |
|  | N | 3 | 3 | 3 | 2 | 3 | 3 | 3 | 3 | 3 |
| 3 | MEAN | 206.0 | 270.3 | 737.3 | 5.53 | 3.217 | 0.687 | 3.87 | 5.43 | 65.3 |
|  | SEM | 74.5 | 84.9 | 178.7 | 1.34 | 0.230 | 0.200 | 0.15 | 0.44 | 4.9 |
|  | N | 3 | 3 | 3 | 3 | 3 | 3 | 3 | 3 | 3 |
| 4 | MEAN | 29.3 | 29.0 | 515.3 | 5.23 | 2.830 | 0.697 | 4.30 | 5.47 | 62.0 |
|  | SEM | 3.3 | 1.0 | 32.7 | 0.95 | 0.263 | 0.163 | 0.17 | 0.78 | 3.5 |
|  | N | 3 | 3 | 3 | 3 | 3 | 3 | 3 | 3 | 3 |
| 5 | MEAN | 40.0 | 30.7 | 358.0 | 3.60 | 3.283 | 0.820 | 4.90 | 5.43 | 69.0 |
|  | SEM | 6.8 | 3.8 | 85.0 | 0.61 | 0.116 | 0.161 | 0.31 | 0.18 | 3.5 |
|  | N | 3 | 3 | 3 | 3 | 3 | 3 | 3 | 3 | 3 |

**Summary of clinical chemistry I – Day 14**

| **GROUP** |  | **AST** | **ALT** | **ALP** | **TBIL** | **CHOL** | **TRIG** | **GLUC** | **UREA** | **CREAT** |
| --- | --- | --- | --- | --- | --- | --- | --- | --- | --- | --- |
| **NO.** |  | **(U/L)** | **(U/L)** | **(U/L)** | **(μmol/L)** | **(mmol/L)** | **(mmol/L)** | **(mmol/L)** | **(mmol/L)** | **(μmol/L)** |
| 1 | MEAN | 25.0 | 34.0 | 529.0 | 2.90 | 3.240 | 0.677 | 5.57 | 6.40 | 59.3 |
|  | SEM | 1.5 | 3.6 | 23.1 | 0.36 | 0.265 | 0.171 | 0.52 | 0.67 | 2.2 |
|  | N | 3 | 3 | 3 | 3 | 3 | 3 | 3 | 3 | 3 |
| 2 | MEAN | 32.7 | 38.3 | 576.3 | 3.50 | 2.697 | 0.873 | 4.70 | 7.07 | 71.3 |
|  | SEM | 2.3 | 1.9 | 54.2 | NA | 0.359 | 0.269 | 0.35 | 0.12 | 6.3 |
|  | N | 3 | 3 | 3 | 1 | 3 | 3 | 3 | 3 | 3 |
| 3 | MEAN | 45.0 | 38.7 | 639.0 | 4.55 | 2.777 | 0.717 | 4.43 | 6.10 | 73.7 |
|  | SEM | 11.8 | 4.2 | 148.7 | 2.05 | 0.192 | 0.134 | 1.04 | 0.50 | 6.7 |
|  | N | 3 | 3 | 3 | 2 | 3 | 3 | 3 | 3 | 3 |
| 4 | MEAN | 27.3 | 30.3 | 542.3 | 2.50 | 2.970 | 0.543 | 4.73 | 5.93 | 61.7 |
|  | SEM | 0.7 | 2.9 | 23.9 | 0.30 | 0.111 | 0.043 | 0.12 | 0.26 | 3.5 |
|  | N | 3 | 3 | 3 | 2 | 3 | 3 | 3 | 3 | 3 |
| 5 | MEAN | 24.3 | 26.3 | 530.0 | NA | 3.277 | 0.480 | 5.13 | 5.67 | 50.7 |
|  | SEM | 3.5 | 2.4 | 165.1 | NA | 0.133 | 0.076 | 0.64 | 0.83 | 1.9 |
|  | N | 3 | 3 | 3 | 0 | 3 | 3 | 3 | 3 | 3 |

**Summary of clinical chemistry I – Day 90**

| **GROUP** |  | **AST** | **ALT** | **ALP** | **TBIL** | **CHOL** | **TRIG** | **GLUC** | **UREA** | **CREAT** |
| --- | --- | --- | --- | --- | --- | --- | --- | --- | --- | --- |
| **NO.** |  | **(U/L)** | **(U/L)** | **(U/L)** | **(μmol/L)** | **(mmol/L)** | **(mmol/L)** | **(mmol/L)** | **(mmol/L)** | **(μmol/L)** |
| 1 | MEAN | 31.0 | 39.0 | 541.3 | 2.50 | 3.007 | 0.733 | 5.10 | 6.27 | 61.0 |
|  | SEM | 1.0 | 12.5 | 39.4 | 0.20 | 0.340 | 0.112 | 0.21 | 0.43 | 3.2 |
|  | N | 3 | 3 | 3 | 2 | 3 | 3 | 3 | 3 | 3 |
| 2 | MEAN | 32.0 | 38.0 | 604.0 | 3.90 | 2.593 | 0.760 | 4.23 | 6.67 | 66.7 |
|  | SEM | 1.5 | 3.5 | 107.9 | NA | 0.367 | 0.156 | 0.32 | 0.61 | 6.1 |
|  | N | 3 | 3 | 3 | 1 | 3 | 3 | 3 | 3 | 3 |
| 3 | MEAN | 39.3 | 41.7 | 709.7 | 5.40 | 2.327 | 0.743 | 4.30 | 5.83 | 66.3 |
|  | SEM | 6.4 | 10.7 | 181.9 | NA | 0.213 | 0.108 | 0.29 | 0.41 | 6.1 |
|  | N | 3 | 3 | 3 | 1 | 3 | 3 | 3 | 3 | 3 |
| 4 | MEAN | 31.0 | 29.7 | 600.7 | NA | 2.800 | 0.810 | 4.83 | 6.33 | 60.0 |
|  | SEM | 1.5 | 0.3 | 19.4 | NA | 0.108 | 0.061 | 0.20 | 0.12 | 1.0 |
|  | N | 3 | 3 | 3 | 0 | 3 | 3 | 3 | 3 | 3 |
| 5 | MEAN | 26.3 | 28.7 | 532.3 | NA | 3.363 | 0.693 | 5.50 | 5.83 | 48.3 |
|  | SEM | 2.3 | 4.8 | 161.4 | NA | 0.175 | 0.088 | 0.26 | 0.55 | 1.2 |
|  | N | 3 | 3 | 3 | 0 | 3 | 3 | 3 | 3 | 3 |

**Summary of clinical chemistry I – Day 129**

| **GROUP** |  | **AST** | **ALT** | **ALP** | **TBIL** | **CHOL** | **TRIG** | **GLUC** | **UREA** | **CREAT** |
| --- | --- | --- | --- | --- | --- | --- | --- | --- | --- | --- |
| **NO.** |  | **(U/L)** | **(U/L)** | **(U/L)** | **(μmol/L)** | **(mmol/L)** | **(mmol/L)** | **(mmol/L)** | **(mmol/L)** | **(μmol/L)** |
| 1 | MEAN | 37.7 | 41.7 | 494.0 | 4.07 | 3.480 | 0.463 | 5.03 | 6.37 | 64.0 |
|  | SEM | 5.4 | 6.7 | 31.2 | 0.48 | 0.428 | 0.050 | 0.59 | 0.72 | 4.2 |
|  | N | 3 | 3 | 3 | 3 | 3 | 3 | 3 | 3 | 3 |
| 2 | MEAN | 38.3 | 33.7 | 592.7 | 3.00 | 2.620 | 0.573 | 4.60 | 6.03 | 73.0 |
|  | SEM | 2.2 | 3.7 | 123.9 | 0.35 | 0.208 | 0.171 | 0.23 | 0.29 | 3.5 |
|  | N | 3 | 3 | 3 | 3 | 3 | 3 | 3 | 3 | 3 |
| 3 | MEAN | 41.3 | 36.0 | 683.0 | 4.90 | 2.650 | 0.630 | 3.77 | 5.67 | 62.0 |
|  | SEM | 9.0 | 7.0 | 137.0 | 2.20 | 0.215 | 0.108 | 0.72 | 0.57 | 3.0 |
|  | N | 3 | 3 | 3 | 2 | 3 | 3 | 3 | 3 | 3 |

**Summary of clinical chemistry I – Day 150**

| **GROUP** |  | **AST** | **ALT** | **ALP** | **TBIL** | **CHOL** | **TRIG** | **GLUC** | **UREA** | **CREAT** |
| --- | --- | --- | --- | --- | --- | --- | --- | --- | --- | --- |
| **NO.** |  | **(U/L)** | **(U/L)** | **(U/L)** | **(μmol/L)** | **(mmol/L)** | **(mmol/L)** | **(mmol/L)** | **(mmol/L)** | **(μmol/L)** |
| 1 | MEAN | 35.7 | 38.7 | 522.7 | 4.45 | 3.133 | 0.470 | 4.87 | 6.60 | 66.0 |
|  | SEM | 6.9 | 7.8 | 32.5 | 0.05 | 0.341 | 0.052 | 0.37 | 0.87 | 4.0 |
|  | N | 3 | 3 | 3 | 2 | 3 | 3 | 3 | 3 | 3 |
| 2 | MEAN | 40.3 | 40.3 | 626.0 | 4.15 | 2.357 | 0.537 | 3.53 | 6.70 | 76.3 |
|  | SEM | 5.5 | 4.1 | 135.3 | 1.75 | 0.209 | 0.067 | 0.24 | 0.17 | 5.8 |
|  | N | 3 | 3 | 3 | 2 | 3 | 3 | 3 | 3 | 3 |
| 3 | MEAN | 38.7 | 38.0 | 704.0 | 4.60 | 2.467 | 0.560 | 3.63 | 6.23 | 68.0 |
|  | SEM | 6.7 | 6.0 | 133.5 | 1.92 | 0.172 | 0.066 | 0.49 | 0.47 | 4.7 |
|  | N | 3 | 3 | 3 | 3 | 3 | 3 | 3 | 3 | 3 |

**Summary of clinical chemistry I – Day 180**

| **GROUP** |  | **AST** | **ALT** | **ALP** | **TBIL** | **CHOL** | **TRIG** | **GLUC** | **UREA** | **CREAT** |
| --- | --- | --- | --- | --- | --- | --- | --- | --- | --- | --- |
| **NO.** |  | **(U/L)** | **(U/L)** | **(U/L)** | **(μmol/L)** | **(mmol/L)** | **(mmol/L)** | **(mmol/L)** | **(mmol/L)** | **(μmol/L)** |
| 1 | MEAN | 27.7 | 33.7 | 462.7 | 2.73 | 3.197 | 0.723 | 5.60 | 6.13 | 66.3 |
|  | SEM | 3.3 | 7.2 | 54.0 | 0.38 | 0.387 | 0.178 | 0.70 | 0.27 | 5.8 |
|  | N | 3 | 3 | 3 | 3 | 3 | 3 | 3 | 3 | 3 |
| 2 | MEAN | 30.3 | 32.3 | 567.3 | 6.10 | 2.480 | 0.723 | 4.53 | 6.97 | 70.0 |
|  | SEM | 4.7 | 0.9 | 126.4 | NA | 0.292 | 0.198 | 0.50 | 0.38 | 8.1 |
|  | N | 3 | 3 | 3 | 1 | 3 | 3 | 3 | 3 | 3 |
| 3 | MEAN | 33.3 | 30.7 | 644.0 | 3.40 | 2.490 | 0.733 | 4.27 | 5.70 | 68.3 |
|  | SEM | 4.9 | 4.9 | 116.2 | 0.95 | 0.171 | 0.055 | 0.27 | 0.21 | 7.4 |
|  | N | 3 | 3 | 3 | 3 | 3 | 3 | 3 | 3 | 3 |

ALP = alkaline phosphatase; ALT = alanine aminotransferase; AST = aspartate aminotransferase; CHOL = cholesterol; CREAT = creatinine; GLUC = glucose; TBIL = total bilirubin.

**Table S5:	Summary of clinical chemistry II – Predose**

| **GROUP** |  | **TPROT** | **ALB** | **GLOB** | **A/G** | **Ca** | **PHOS** | **Na** | **K** | **Cl** |
| --- | --- | --- | --- | --- | --- | --- | --- | --- | --- | --- |
| **NO.** |  | **(g/L)** | **(g/L)** | **(g/L)** |  | **(mmol/L)** | **(mmol/L)** | **(mmol/L)** | **(mmol/L)** | **(mmol/L)** |
| 1 | MEAN | 70.80 | 49.00 | 21.80 | 2.250 | 2.577 | 1.753 | 145.0 | 4.150 | 102.2 |
|  | SEM | 0.44 | 0.30 | 0.50 | 0.061 | 0.003 | 0.167 | 0.6 | 0.178 | 2.0 |
|  | N | 3 | 3 | 3 | 3 | 3 | 3 | 3 | 3 | 3 |
| 2 | MEAN | 74.40 | 49.13 | 25.27 | 1.947 | 2.577 | 2.050 | 147.7 | 4.033 | 101.4 |
|  | SEM | 1.37 | 1.09 | 0.71 | 0.064 | 0.046 | 0.202 | 2.3 | 0.240 | 1.7 |
|  | N | 3 | 3 | 3 | 3 | 3 | 3 | 3 | 3 | 3 |
| 3 | MEAN | 77.03 | 51.23 | 25.80 | 2.013 | 2.667 | 1.783 | 147.3 | 3.983 | 104.0 |
|  | SEM | 1.84 | 0.38 | 2.14 | 0.176 | 0.009 | 0.085 | 1.3 | 0.117 | 1.1 |
|  | N | 3 | 3 | 3 | 3 | 3 | 3 | 3 | 3 | 3 |
| 4 | MEAN | 72.27 | 48.43 | 23.83 | 2.060 | 2.573 | 1.773 | 148.0 | 3.873 | 103.8 |
|  | SEM | 2.18 | 0.44 | 2.09 | 0.165 | 0.015 | 0.165 | 0.6 | 0.107 | 2.1 |
|  | N | 3 | 3 | 3 | 3 | 3 | 3 | 3 | 3 | 3 |
| 5 | MEAN | 72.63 | 47.87 | 24.77 | 1.933 | 2.527 | 1.930 | 145.3 | 3.760 | 103.2 |
|  | SEM | 0.90 | 0.13 | 0.77 | 0.056 | 0.035 | 0.113 | 0.3 | 0.104 | 1.1 |
|  | N | 3 | 3 | 3 | 3 | 3 | 3 | 3 | 3 | 3 |

**Summary of clinical chemistry II – Day 3**

| **GROUP** |  | **TPROT** | **ALB** | **GLOB** | **A/G** | **Ca** | **PHOS** | **Na** | **K** | **Cl** |
| --- | --- | --- | --- | --- | --- | --- | --- | --- | --- | --- |
| **NO.** |  | **(g/L)** | **(g/L)** | **(g/L)** |  | **(mmol/L)** | **(mmol/L)** | **(mmol/L)** | **(mmol/L)** | **(mmol/L)** |
| 1 | MEAN | 71.43 | 49.23 | 22.20 | 2.223 | 2.560 | 1.723 | 147.3 | 4.140 | 105.8 |
|  | SEM | 1.10 | 0.34 | 0.76 | 0.058 | 0.021 | 0.154 | 0.9 | 0.032 | 0.9 |
|  | N | 3 | 3 | 3 | 3 | 3 | 3 | 3 | 3 | 3 |
| 2 | MEAN | 74.20 | 46.37 | 27.83 | 1.700 | 2.547 | 1.617 | 147.0 | 3.817 | 103.6 |
|  | SEM | 0.55 | 1.78 | 2.33 | 0.190 | 0.065 | 0.203 | 3.0 | 0.236 | 2.2 |
|  | N | 3 | 3 | 3 | 3 | 3 | 3 | 3 | 3 | 3 |
| 3 | MEAN | 74.00 | 47.40 | 26.60 | 1.790 | 2.530 | 1.653 | 149.0 | 3.900 | 105.6 |
|  | SEM | 1.59 | 0.87 | 1.20 | 0.085 | 0.056 | 0.170 | 1.5 | 0.193 | 0.6 |
|  | N | 3 | 3 | 3 | 3 | 3 | 3 | 3 | 3 | 3 |
| 4 | MEAN | 67.30 | 38.27 | 29.03 | 1.330 | 2.350 | 1.277 | 142.3 | 3.697 | 103.3 |
|  | SEM | 3.39 | 0.97 | 2.42 | 0.074 | 0.015 | 0.145 | 0.7 | 0.109 | 1.2 |
|  | N | 3 | 3 | 3 | 3 | 3 | 3 | 3 | 3 | 3 |
| 5 | MEAN | 66.93 | 36.03 | 30.90 | 1.170 | 2.383 | 1.237 | 145.3 | 3.920 | 100.3 |
|  | SEM | 0.24 | 0.18 | 0.42 | 0.021 | 0.067 | 0.074 | 2.4 | 0.197 | 1.5 |
|  | N | 3 | 3 | 3 | 3 | 3 | 3 | 3 | 3 | 3 |

**Summary of clinical chemistry II – Day 14**

| **GROUP** |  | **TPROT** | **ALB** | **GLOB** | **A/G** | **Ca** | **PHOS** | **Na** | **K** | **Cl** |
| --- | --- | --- | --- | --- | --- | --- | --- | --- | --- | --- |
| **NO.** |  | **(g/L)** | **(g/L)** | **(g/L)** |  | **(mmol/L)** | **(mmol/L)** | **(mmol/L)** | **(mmol/L)** | **(mmol/L)** |
| 1 | MEAN | 70.67 | 48.37 | 22.30 | 2.177 | 2.627 | 1.837 | 147.3 | 4.393 | 104.9 |
|  | SEM | 0.69 | 0.03 | 0.70 | 0.072 | 0.022 | 0.167 | 1.7 | 0.104 | 1.5 |
|  | N | 3 | 3 | 3 | 3 | 3 | 3 | 3 | 3 | 3 |
| 2 | MEAN | 74.73 | 47.60 | 27.13 | 1.763 | 2.647 | 1.980 | 149.3 | 4.433 | 104.8 |
|  | SEM | 1.67 | 1.44 | 1.36 | 0.113 | 0.033 | 0.143 | 1.5 | 0.275 | 0.6 |
|  | N | 3 | 3 | 3 | 3 | 3 | 3 | 3 | 3 | 3 |
| 3 | MEAN | 74.03 | 48.13 | 25.90 | 1.890 | 2.670 | 1.667 | 151.3 | 4.397 | 105.8 |
|  | SEM | 2.35 | 0.73 | 2.23 | 0.176 | 0.038 | 0.240 | 1.3 | 0.423 | 1.5 |
|  | N | 3 | 3 | 3 | 3 | 3 | 3 | 3 | 3 | 3 |
| 4 | MEAN | 70.60 | 44.67 | 25.93 | 1.743 | 2.547 | 1.830 | 149.7 | 4.177 | 104.8 |
|  | SEM | 1.89 | 0.09 | 1.90 | 0.124 | 0.049 | 0.130 | 1.5 | 0.037 | 0.3 |
|  | N | 3 | 3 | 3 | 3 | 3 | 3 | 3 | 3 | 3 |
| 5 | MEAN | 69.27 | 39.20 | 30.07 | 1.307 | 2.463 | 1.677 | 149.3 | 3.930 | 105.3 |
|  | SEM | 0.54 | 1.25 | 0.93 | 0.083 | 0.032 | 0.097 | 1.2 | 0.156 | 0.6 |
|  | N | 3 | 3 | 3 | 3 | 3 | 3 | 3 | 3 | 3 |

**Summary of clinical chemistry II – Day 90**

| **GROUP** |  | **TPROT** | **ALB** | **GLOB** | **A/G** | **Ca** | **PHOS** | **Na** | **K** | **Cl** |
| --- | --- | --- | --- | --- | --- | --- | --- | --- | --- | --- |
| **NO.** |  | **(g/L)** | **(g/L)** | **(g/L)** |  | **(mmol/L)** | **(mmol/L)** | **(mmol/L)** | **(mmol/L)** | **(mmol/L)** |
| 1 | MEAN | 73.87 | 47.77 | 26.10 | 1.837 | 2.557 | 1.600 | 148.0 | 3.700 | 103.9 |
|  | SEM | 1.17 | 0.61 | 0.83 | 0.054 | 0.043 | 0.219 | 1.5 | 0.111 | 0.9 |
|  | N | 3 | 3 | 3 | 3 | 3 | 3 | 3 | 3 | 3 |
| 2 | MEAN | 75.67 | 47.80 | 27.87 | 1.717 | 2.600 | 1.647 | 147.0 | 3.800 | 104.5 |
|  | SEM | 0.99 | 0.68 | 0.34 | 0.012 | 0.046 | 0.215 | 2.1 | 0.185 | 1.3 |
|  | N | 3 | 3 | 3 | 3 | 3 | 3 | 3 | 3 | 3 |
| 3 | MEAN | 77.87 | 49.33 | 28.53 | 1.753 | 2.630 | 1.393 | 151.0 | 3.683 | 105.9 |
|  | SEM | 1.87 | 0.43 | 2.30 | 0.162 | 0.025 | 0.174 | 1.5 | 0.268 | 1.4 |
|  | N | 3 | 3 | 3 | 3 | 3 | 3 | 3 | 3 | 3 |
| 4 | MEAN | 73.57 | 47.07 | 26.50 | 1.807 | 2.530 | 1.607 | 147.3 | 3.967 | 103.4 |
|  | SEM | 2.22 | 1.22 | 2.38 | 0.176 | 0.040 | 0.108 | 0.7 | 0.265 | 0.5 |
|  | N | 3 | 3 | 3 | 3 | 3 | 3 | 3 | 3 | 3 |
| 5 | MEAN | 72.77 | 47.43 | 25.33 | 1.873 | 2.523 | 1.453 | 146.3 | 3.870 | 103.4 |
|  | SEM | 0.74 | 0.45 | 0.60 | 0.047 | 0.078 | 0.196 | 0.3 | 0.385 | 0.8 |
|  | N | 3 | 3 | 3 | 3 | 3 | 3 | 3 | 3 | 3 |

**Summary of clinical chemistry II – Day 129**

| **GROUP** |  | **TPROT** | **ALB** | **GLOB** | **A/G** | **Ca** | **PHOS** | **Na** | **K** | **Cl** |
| --- | --- | --- | --- | --- | --- | --- | --- | --- | --- | --- |
| **NO.** |  | **(g/L)** | **(g/L)** | **(g/L)** |  | **(mmol/L)** | **(mmol/L)** | **(mmol/L)** | **(mmol/L)** | **(mmol/L)** |
| 1 | MEAN | 72.07 | 48.67 | 23.40 | 2.083 | 2.547 | 1.640 | 147.3 | 4.080 | 104.4 |
|  | SEM | 1.11 | 0.41 | 0.72 | 0.047 | 0.012 | 0.188 | 0.3 | 0.070 | 1.1 |
|  | N | 3 | 3 | 3 | 3 | 3 | 3 | 3 | 3 | 3 |
| 2 | MEAN | 73.07 | 47.87 | 25.20 | 1.900 | 2.573 | 1.843 | 148.0 | 3.947 | 103.1 |
|  | SEM | 1.27 | 0.88 | 0.42 | 0.015 | 0.012 | 0.078 | 1.0 | 0.187 | 1.6 |
|  | N | 3 | 3 | 3 | 3 | 3 | 3 | 3 | 3 | 3 |
| 3 | MEAN | 74.37 | 48.87 | 25.50 | 1.943 | 2.630 | 1.603 | 149.0 | 4.133 | 105.0 |
|  | SEM | 2.28 | 1.36 | 2.01 | 0.175 | 0.040 | 0.178 | 0.6 | 0.070 | 0.9 |
|  | N | 3 | 3 | 3 | 3 | 3 | 3 | 3 | 3 | 3 |

**Summary of clinical chemistry II – Day 150**

| **GROUP** |  | **TPROT** | **ALB** | **GLOB** | **A/G** | **Ca** | **PHOS** | **Na** | **K** | **Cl** |
| --- | --- | --- | --- | --- | --- | --- | --- | --- | --- | --- |
| **NO.** |  | **(g/L)** | **(g/L)** | **(g/L)** |  | **(mmol/L)** | **(mmol/L)** | **(mmol/L)** | **(mmol/L)** | **(mmol/L)** |
| 1 | MEAN | 72.97 | 51.50 | 21.47 | 2.407 | 2.550 | 1.780 | 144.0 | 3.917 | 102.0 |
|  | SEM | 1.11 | 0.59 | 0.93 | 0.104 | 0.017 | 0.101 | 1.5 | 0.083 | 1.9 |
|  | N | 3 | 3 | 3 | 3 | 3 | 3 | 3 | 3 | 3 |
| 2 | MEAN | 72.60 | 48.73 | 23.87 | 2.043 | 2.593 | 1.853 | 146.7 | 3.910 | 101.7 |
|  | SEM | 1.24 | 1.42 | 0.22 | 0.076 | 0.018 | 0.152 | 0.9 | 0.208 | 0.7 |
|  | N | 3 | 3 | 3 | 3 | 3 | 3 | 3 | 3 | 3 |
| 3 | MEAN | 76.60 | 51.73 | 24.87 | 2.093 | 2.627 | 1.540 | 146.7 | 3.630 | 103.4 |
|  | SEM | 1.23 | 0.86 | 1.13 | 0.115 | 0.037 | 0.166 | 0.3 | 0.145 | 0.8 |
|  | N | 3 | 3 | 3 | 3 | 3 | 3 | 3 | 3 | 3 |

**Summary of clinical chemistry II – Day 180**

| **GROUP** |  | **TPROT** | **ALB** | **GLOB** | **A/G** | **Ca** | **PHOS** | **Na** | **K** | **Cl** |
| --- | --- | --- | --- | --- | --- | --- | --- | --- | --- | --- |
| **NO.** |  | **(g/L)** | **(g/L)** | **(g/L)** |  | **(mmol/L)** | **(mmol/L)** | **(mmol/L)** | **(mmol/L)** | **(mmol/L)** |
| 1 | MEAN | 72.00 | 46.90 | 25.10 | 1.883 | 2.477 | 1.680 | 149.0 | 4.283 | 106.2 |
|  | SEM | 2.06 | 1.14 | 1.50 | 0.119 | 0.026 | 0.180 | 3.2 | 0.064 | 2.4 |
|  | N | 3 | 3 | 3 | 3 | 3 | 3 | 3 | 3 | 3 |
| 2 | MEAN | 70.47 | 46.33 | 24.13 | 1.930 | 2.530 | 1.920 | 148.7 | 4.117 | 104.3 |
|  | SEM | 1.02 | 1.69 | 1.01 | 0.144 | 0.066 | 0.147 | 2.4 | 0.149 | 1.5 |
|  | N | 3 | 3 | 3 | 3 | 3 | 3 | 3 | 3 | 3 |
| 3 | MEAN | 74.97 | 49.97 | 25.00 | 2.017 | 2.660 | 1.467 | 147.0 | 3.933 | 103.2 |
|  | SEM | 1.11 | 0.94 | 1.51 | 0.154 | 0.035 | 0.191 | 0.6 | 0.207 | 0.7 |
|  | N | 3 | 3 | 3 | 3 | 3 | 3 | 3 | 3 | 3 |

A/G = albumin/globulin ratio; ALB = albumin; Ca = calcium; Cl = chloride; GLOB = globulin; K = potassium; Na = sodium; PHOS = phosphorus; TPROT = total protein.

#### **Summary of coagulation**

**Table S3:**	**Summary of coagulation – Predose**

| **GROUP NO.** |  | **PT**  **(Sec)** | **APTT**  **(Sec)** | **FIB**  **(g/L)** |
| --- | --- | --- | --- | --- |
| 1 | MEAN | 13.00 | 39.60 | 1.85 |
|  | SEM | 0.25 | 4.47 | 0.21 |
|  | N | 3 | 3 | 3 |
| 2 | MEAN | 12.90 | 33.47 | 2.20 |
|  | SEM | 0.44 | 0.83 | 0.15 |
|  | N | 3 | 3 | 3 |
| 3 | MEAN | 12.53 | 30.13 | 2.21 |
|  | SEM | 0.19 | 3.74 | 0.11 |
|  | N | 3 | 3 | 3 |
| 4 | MEAN | 13.03 | 32.03 | 2.12 |
|  | SEM | 0.30 | 1.49 | 0.12 |
|  | N | 3 | 3 | 3 |
| 5 | MEAN | 12.80 | 28.00 | 2.09 |
|  | SEM | 0.25 | 2.05 | 0.27 |
|  | N | 3 | 3 | 3 |

**Summary of coagulation – Day 3**

| **GROUP NO.** |  | **PT**  **(Sec)** | **APTT**  **(Sec)** | **FIB**  **(g/L)** |
| --- | --- | --- | --- | --- |
| 1 | MEAN | 13.27 | 37.77 | 2.15 |
|  | SEM | 0.23 | 5.44 | 0.11 |
|  | N | 3 | 3 | 3 |
| 2 | MEAN | 12.53 | 31.57 | 3.17 |
|  | SEM | 0.44 | 0.46 | 0.31 |
|  | N | 3 | 3 | 3 |
| 3 | MEAN | 15.00 | 28.53 | 2.30 |
|  | SEM | 0.85 | 2.60 | 0.42 |
|  | N | 3 | 3 | 3 |
| 4 | MEAN | 13.33 | 33.83 | 6.85 |
|  | SEM | 0.33 | 2.20 | 0.50 |
|  | N | 3 | 3 | 3 |
| 5 | MEAN | 13.23 | 34.30 | 7.59 |
|  | SEM | 0.38 | 4.51 | 0.83 |
|  | N | 3 | 3 | 3 |

**Summary of coagulation – Day 14**

| **GROUP NO.** |  | **PT**  **(Sec)** | **APTT**  **(Sec)** | **FIB**  **(g/L)** |
| --- | --- | --- | --- | --- |
| 1 | MEAN | 13.50 | 34.57 | 1.83 |
|  | SEM | 0.32 | 4.60 | 0.15 |
|  | N | 3 | 3 | 3 |
| 2 | MEAN | 12.77 | 30.73 | 2.39 |
|  | SEM | 0.35 | 1.23 | 0.17 |
|  | N | 3 | 3 | 3 |
| 3 | MEAN | 12.83 | 28.43 | 2.26 |
|  | SEM | 0.18 | 2.94 | 0.08 |
|  | N | 3 | 3 | 3 |
| 4 | MEAN | 13.63 | 31.93 | 2.29 |
|  | SEM | 0.24 | 1.79 | 0.04 |
|  | N | 3 | 3 | 3 |
| 5 | MEAN | 12.60 | 26.73 | 2.36 |
|  | SEM | 0.21 | 2.98 | 0.22 |
|  | N | 3 | 3 | 3 |

**Summary of coagulation – Day 90**

| **GROUP NO.** |  | **PT**  **(Sec)** | **APTT**  **(Sec)** | **FIB**  **(g/L)** |
| --- | --- | --- | --- | --- |
| 1 | MEAN | 13.83 | 34.07 | 2.03 |
|  | SEM | 0.22 | 2.60 | 0.28 |
|  | N | 3 | 3 | 3 |
| 2 | MEAN | 13.03 | 30.00 | 2.15 |
|  | SEM | 0.22 | 0.38 | 0.07 |
|  | N | 3 | 3 | 3 |
| 3 | MEAN | 13.07 | 29.30 | 2.17 |
|  | SEM | 0.32 | 2.66 | 0.10 |
|  | N | 3 | 3 | 3 |
| 4 | MEAN | 13.83 | 30.17 | 2.04 |
|  | SEM | 0.24 | 0.90 | 0.11 |
|  | N | 3 | 3 | 3 |
| 5 | MEAN | 13.43 | 29.53 | 2.00 |
|  | SEM | 0.18 | 3.78 | 0.32 |
|  | N | 3 | 3 | 3 |

**Summary of coagulation – Day 129**

| **GROUP NO.** |  | **PT**  **(Sec)** | **APTT**  **(Sec)** | **FIB**  **(g/L)** |
| --- | --- | --- | --- | --- |
| 1 | MEAN | 13.63 | 34.63 | 2.04 |
|  | SEM | 0.27 | 4.01 | 0.15 |
|  | N | 3 | 3 | 3 |
| 2 | MEAN | 13.50 | 30.70 | 2.79 |
|  | SEM | 0.40 | 1.37 | 0.46 |
|  | N | 3 | 3 | 3 |
| 3 | MEAN | 13.03 | 30.43 | 2.24 |
|  | SEM | 0.32 | 2.38 | 0.24 |
|  | N | 3 | 3 | 3 |

**Summary of coagulation – Day 150**

| **GROUP NO.** |  | **PT**  **(Sec)** | **APTT**  **(Sec)** | **FIB**  **(g/L)** |
| --- | --- | --- | --- | --- |
| 1 | MEAN | 13.53 | 32.10 | 1.87 |
|  | SEM | 0.32 | 2.76 | 0.20 |
|  | N | 3 | 3 | 3 |
| 2 | MEAN | 13.67 | 31.63 | 2.18 |
|  | SEM | 0.45 | 0.59 | 0.06 |
|  | N | 3 | 3 | 3 |
| 3 | MEAN | 13.40 | 29.23 | 2.13 |
|  | SEM | 0.15 | 1.59 | 0.11 |
|  | N | 3 | 3 | 3 |

**Summary of coagulation – Day 180**

| **GROUP NO.** |  | **PT**  **(Sec)** | **APTT**  **(Sec)** | **FIB**  **(g/L)** |
| --- | --- | --- | --- | --- |
| 1 | MEAN | 13.63 | 36.40 | 1.94 |
|  | SEM | 0.17 | 3.90 | 0.21 |
|  | N | 3 | 3 | 3 |
| 2 | MEAN | 12.87 | 31.43 | 2.35 |
|  | SEM | 0.28 | 1.41 | 0.09 |
|  | N | 3 | 3 | 3 |
| 3 | MEAN | 13.10 | 28.70 | 2.26 |
|  | SEM | 0.10 | 2.31 | 0.13 |
|  | N | 3 | 3 | 3 |

APTT = activated partial thromboplastin time; FIB = fibrinogen; PT = prothrombin.

#### **Allometric exponents for Fc-engineered mAbs and RibobNAbs, as well as NHP and human PK parameters**

Table S4:	Allometric exponents (b) for Fc-engineered mAbs^79^ and RibobNAbs, as well as NHP and human PK parameters

| Parameter | Exponent NHP | NHP parameters | Predicted human parameters |
| --- | --- | --- | --- |
| V_c_ | 0.95 | 2.32 (L) | 2.03 (L) |
| V_p_ | 0.95 | 1.77 (L) | 1.55 (L) |
| CL | 0.55 | 0.13 (L/day) | 0.04 (L/day) |
| Q | 0.6 | 0.74 (L/day) | 0.25 (L/day) |
| k_a_* | 1 |  |  |
| F* | 1 |  |  |

Abbreviations: Fc = fragment crystallizable; mAb = monoclonal antibody; NHP = non-human primate; V_c_ = volume of distribution in the central compartment; V_p_ = volume of distribution in the peripheral compartment; CL = clearance from the central compartment; Q = intercompartmental clearance between central and peripheral compartment, *RibobNAb specific parameters: k_a_= translational rate, F= translational efficiency.
